# An engineered nanobody inhibitor for molecular-to-circuit control of opioid receptor function

**DOI:** 10.64898/2026.01.12.698943

**Authors:** Zoé Valbret, Paul J. Lamothe-Molina, Sofia Papadogkonaki, Antoine Koehl, Jan Vincent V. Arafiles, Ziva Shapiro Tuchman, Nicole M. Fisher, Arthur Radoux-Mergault, Annika Canziani, Gaoyang Huang, Mark von Zastrow, Mickey Kosloff, Christian P. R. Hackenberger, Aashish Manglik, Tommaso Patriarchi, Miriam Stoeber

**Author notes:** co-first authors (Zoé Valbret, Paul J. Lamothe-Molina).

## Abstract

Opioid receptors (ORs) orchestrate pain relief, reward, and dependence, yet their signaling arises from diverse cell types and subcellular compartments that cannot be selectively interrogated with existing pharmacological or genetic approaches. Single-domain antibodies, or nanobodies (Nbs), can probe receptor states, but their potential as tools for controlling native receptor signaling at the system level has remained unexplored. Here, we engineer a suite of high-affinity intracellular Nbs that bind active ORs through structure-guided evolution and in silico design. Iterative optimization yields Nb64, a potent inhibitor that rapidly suppresses transducer engagement, receptor internalization, and downstream signaling, including endogenous pathways in neuronal cells. Organelle targeting highlights Nb64’s capacity to control OR activity with subcellular precision, while bio-reversible cell-penetrating peptide (CPP) conjugation enables non-genetic cytosolic delivery. Cell-type-specific expression of Nb64 in VTA interneurons attenuates fentanyl-evoked dopamine release and behavioral responses in mice, demonstrating targeted control of opioid actions in vivo. Nb64 provides a versatile strategy for dissecting OR biology and establishes a generalizable framework for precision inhibition of native GPCR signaling in vivo.

## Introduction

Neuromodulatory G protein-coupled receptors (GPCRs) regulate neuronal activity and synaptic transmission through intracellular signaling cascades, thereby shaping brain function and animal behavior. Although decades of research have elucidated the molecular principles of GPCR signaling and drug action, it has also become clear that receptor function is highly context-dependent, varying across cell types, neural circuits, and subcellular compartments ^1–5^. However, these dimensions of GPCR signaling remain difficult to interrogate experimentally due to the lack of molecularly precise tools capable of intervening selectively at the level of endogenous receptors. As a result, linking the activity of specific GPCR populations to physiological and behavioral outputs remains a central challenge at the interface of chemical biology and translational neuroscience.

Opioid receptors (ORs), including μOR, δOR, κOR, and nociceptin NOP-R, are neuromodulatory GPCRs that regulate pain, reward, and mood. They are targets of endogenous opioid neuropeptides and clinically important opioid drugs. ORs mediate both the analgesic actions of opioid drugs and their deleterious effects, including respiratory depression, tolerance, and dependence, motivating detailed investigation of their mechanisms of action. ORs are widely expressed across the nervous system, including nociceptive, affective/reward, and brainstem respiratory circuits. However, the contributions of specific cellular and subcellular receptor pools to analgesia, respiratory depression, and reinforcement remain incompletely understood ^6–8^. The challenge is compounded by the fact that agonists activate ORs in different subcellular compartments, including post- and presynaptic terminals as well as intracellular organelles, where they can engage different transducers and generate spatially segregated signaling events ^9–11^. These compartmentalized OR pools may mediate unique responses to neuropeptides and opioid drugs that cannot yet be experimentally resolved.

Traditional pharmacological agents lack the spatial specificity needed to disentangle distinct OR functions, as all receptor-expressing cells and subcellular receptor pools are modulated simultaneously. Genetic strategies, such as conditional knockouts ^12–14^, provide cell-type resolution but can elicit compensatory adaptations and do not permit analysis of receptor function with subcellular precision. Other approaches, including photoswitchable ligands, can achieve receptor modulation with spatial resolution ^15–17^, however, they require optical fiber implantation and depend on local ligand delivery and diffusion, limiting their broader applicability.

Nanobodies (Nbs), single-domain antibody fragments that target conformational epitopes with high selectivity, have emerged as powerful genetically encoded tools to interrogate or modulate GPCRs in living cells ^18–20^. Active-state GPCR-binding Nbs have enabled visualization of ligand-specific receptor activation and, in some cases, selective modulation of GPCR signaling at distinct subcellular sites ^21–23^. However, existing OR-targeting Nbs that bind to the intracellular receptor face, such as Nb39 and Nb33 ^24^, function primarily as biosensors and do not inhibit OR signaling ^23,25^, because their affinities are insufficient to outcompete endogenous transducers.

To overcome these limitations, we engineered high-affinity intracellular Nbs that selectively bind agonist-bound ORs and inhibit downstream signaling. Using structure-guided engineering and functional screening, we generated active-state-dependent Nb variants that exhibit improved receptor engagement. The Nbs sterically prevent coupling to G proteins, GRKs, and β-arrestins, and inhibit μOR-mediated signal transduction and receptor internalization across a wide range of expression levels. We further created a Golgi-targeted Nb that selectively blocks δOR activation at this intracellular organelle, providing a path toward dissecting compartment-specific receptor activity.

Finally, we established the utility of the Nb blocker in vivo by expressing it in μOR-positive GABAergic neurons of the ventral tegmental area (VTA). By preventing μOR-mediated disinhibition of dopamine neurons, this Nb attenuated fentanyl-evoked nucleus accumbens (NAc) dopamine release and reduced opioid-induced behavioral responses in awake, behaving mice.

Together, the findings introduce a nanobody-based strategy for precision inhibition of OR signaling through intracellular competitive binding rather than ligand delivery or receptor knock-out. By enabling molecular, cellular, and subcellular specificity, this approach links receptor-level mechanisms to circuit-level function, allowing causal interrogation of how discrete OR pools shape physiology and behavior and enabling spatially precise modulation of native GPCR pathways in vivo.

## Results

### Affinity maturation yields high-affinity nanobodies for active-state mu-opioid receptor

The OR-targeting nanobodies Nb33 and Nb39, which are nearly identical in sequence (Suppl. Fig. S1A) bind agonist-occupied opioid receptors (μOR, δOR, κOR) without disrupting downstream signaling and have served as conformational biosensors in living cells ^23,26,27^. The high-resolution structure of the agonist-bound, Nb39-stabilized μOR showed that Nb39 interacts with the receptor’s intracellular face through its variable complementarity-determining regions (CDR) 2 and 3 and hydrogen bonds formed by the third framework region ^24^. Because the Nb39–μOR interface overlaps with the G protein-binding site ^28^, we hypothesized that affinity-matured Nb39 variants may sterically block transducer engagement and thereby inhibit OR signaling.

To develop high-affinity variants, we constructed a library of Nb39 mutants containing conservative amino acid substitutions in CDR2 and in the framework region of the receptor-binding surface (Fig. 1A, Suppl. Fig. S1B). The Nb library (∼1.5 × 10⁹ theoretical diversity) was displayed on the surface of yeast and subjected to successive rounds of successive rounds of magnetic cell separation (MACS), retaining Nbs that bind fluorescently-labeled, agonist (BU72)-bound μOR (Fig. 1B). Receptor concentration was progressively decreased to enrich higher-affinity Nb clones, followed by a kinetic selection step favoring slowly dissociating Nbs. After the final selection round, this time using fluorescence-activated cell sorting (FACS), seven distinct clones carrying mutations in both CDR2 and the framework were recovered (Fig. 1C, Suppl. Fig. S1B). The Nb variants exhibited a 100-fold improvement in apparent affinity compared to the original Nb39 in on-yeast titrations (Fig. 1D, Suppl. Fig. S1C). Clone 2, termed Nb62, was selected for further characterization as it contained the consensus sequence of the high-affinity Nbs.

**Figure 1.**
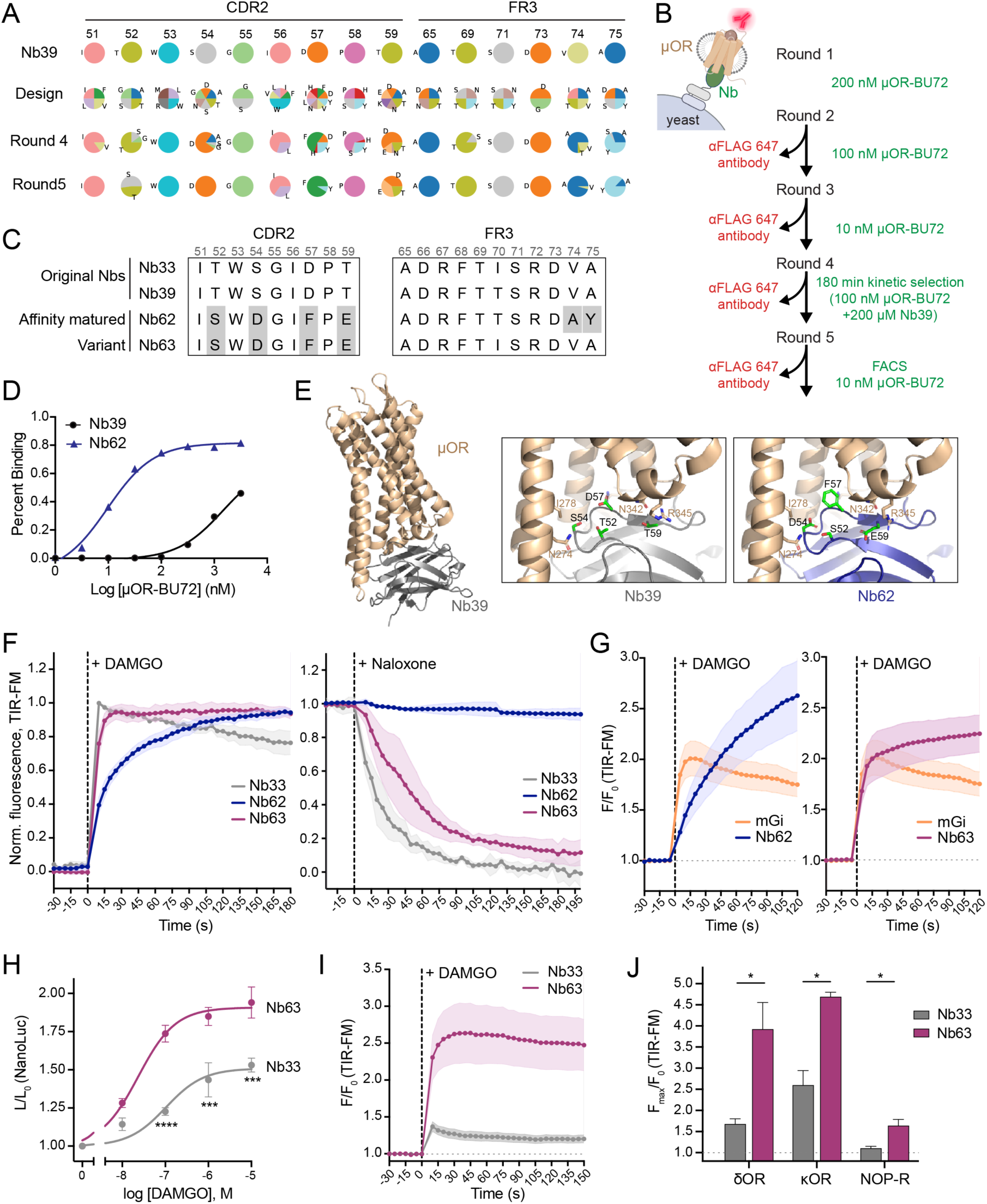
Development of a high-affinity nanobody for active-state opioid receptors. (A) Nb39-based nanobody library containing substitutions in CDR2 and framework region 3. Variant enrichment after selection rounds 4 and 5 based on deep sequencing is shown. (B) Schematic representation of the yeast display screening approach used to isolate high-affinity, active-state μOR-selective Nbs. Purified μOR, bound to the agonist BU72 and fluorescently labeled with AF647-conjugated anti-FLAG antibody, was used. (C) CDR2 and framework region 3 (FR3) sequences of the original (Nb33, Nb39) and affinity-matured (Nb62, Nb63) nanobodies are shown. Shaded positions indicate amino acid substitutions relative to Nb39. (D) On-yeast titration of Nb39- or Nb62-expressing yeast with BU72-bound μOR-AF647 yields EC50 values of 1.5 μM and 10 nM, respectively. (E) Left: structure of the μOR-Nb39 complex (PDB 5C1M). Middle and right: CDR2 residues substituted in Nb62 (blue), relative to Nb39 (grey), shown as green sticks. μOR residues that may interact with Nb62-specific residues are shown as wheat-colored sticks. (F) Left: recruitment kinetics of EGFP-Nbs to μOR upon addition of the agonist DAMGO (1 μM). N = 3. Right: detachment of EGFP-Nbs from μOR after addition of the antagonist naloxone (10 μM; cells pre-treated with 1 μM DAMGO). N = 3. Quantification of EGFP signal in live-cell TIR-FM time-series recordings (5 s interval) of HEK293 cells expressing μOR and EGFP-Nb33, -Nb62, or -Nb63. EGFP intensities were normalized between 0 (no agonist) and 1 (1 μM DAMGO). Mean +/- SEM. (G) Recruitment kinetics detected in TIR-FM time-series recordings (5 s interval) of EGFP-Nb62 or EGFP-Nb63 and mRuby-mGi1 to μOR in HEK293 cells. DAMGO (10 μM) was added. F0 represents the average fluorescence intensity before agonist addition. N = 3, mean +/- SEM. (H) NanoLuc assay measuring μOR interaction with Nb33 or Nb63 based on complementation of Lyn-SmBiT at the plasma membrane and LgBiT-Nb. Concentration-dependent engagement of LgBiT-Nb33 or LgBiT-Nb63 with μOR. Luminescence was normalized to the untreated control cells and to the signal before agonist addition. N = 3, mean +/- SEM. Regression curves were fitted with a Hill slope of 1. ***P < 0.001, ****P < 0.0001, two-way ANOVA with Sidak’s multiple-comparisons test. (I) Recruitment of Nb33 and Nb63 in μOR-expressing HEK293 cells co-expressing both nanobodies as detected in TIR-FM time-series recordings (5 s interval). DAMGO (10 μM) was added. F0 represents the average fluorescence intensity before agonist addition. N = 7, mean +/- SEM. (J) Maximum fold-change in signal detected during TIR-FM time-lapse recordings of EGFP-Nb33 or EGFP-Nb63 binding to δOR, κOR, or NOP-R upon addition of 10 μM peptide agonists DPDPE, DynA, or NOP, respectively. N = 3, mean +/- SEM. ***P < 0.05, Welch’s t-test.

Nb62 differs from Nb39 by six amino acids, four within CDR2 and two in the third framework region (Fig. 1C). To understand how these substitutions enhance μOR binding, we modeled Nb62 using the Nb39 structure as a template (Fig. 1E, see methods). Three mutations in CDR2 were predicted to create new contacts with μOR (Suppl. Fig. S1D): S54D^Nb^ forms a hydrogen bond with N274^μOR^, D57F^Nb^ introduces nonpolar interactions with I278^μOR^, and T59E^Nb^ forms a salt bridge with R345^μOR^. Additionally, A75Y^Nb^ in the third framework region may form hydrogen bonds with S261^μOR^ or R258^μOR^ (Suppl. Fig. S1D).

We next tested whether the affinity gains achieved by *in vitro* selection translated to functional receptor engagement in the cellular context.

### Nanobody optimization for rapid, high-affinity binding of opioid receptors in living cells

EGFP-tagged Nb62 was expressed in HEK293 cells stably expressing μOR (HEK-μOR), and its recruitment to receptors in the plasma membrane was visualized by total internal reflection fluorescence microscopy (TIR-FM) time series. Activation of μOR by the peptide agonist DAMGO led to Nb62 recruitment, but with a slow onset over several minutes (Fig. 1F). This kinetic profile differed from the rapid recruitment of the original Nb33. Moreover, Nb62 dissociated slowly and incompletely from μOR after adding the antagonist naloxone, unlike Nb33 (Fig. 1F).

Since our goal was to develop a Nb capable of blocking off transducers, including G proteins that exhibit rapid receptor interactions, the slow recruitment of Nb62 was suboptimal. A side-by-side comparison of Nb62 and the Gi protein probe miniGi confirmed that Gi binding to agonist-bound μOR was considerably faster, suggesting that Nb62 was not suited to interfere with fast transducer engagement (Fig. 1G).

Because rapid competition with signaling partners of the μOR requires both high affinity and appropriate conformational flexibility of the paratope, we reasoned that the framework substitutions in Nb62, while affinity-enhancing, may inadvertently affect the conformational flexibility required for fast receptor engagement. To improve μOR engagement kinetics, we therefore engineered Nb63, which retained the four CDR2 substitutions of Nb62 but restored the original framework sequence (Fig. 1C). Indeed, Nb63 exhibited markedly faster kinetics in the TIR-FM recruitment assay: it rapidly bound to activated μOR and dissociated quickly upon antagonist addition (Fig.1F). Furthermore, Nb63 and miniGi recruitment to μOR occurred with similar kinetics (Fig. 1G).

To test whether Nb63 retains enhanced binding relative to the original Nbs, we quantified the agonist-driven recruitment of Nb63 and Nb33 to μOR using a split nanoluciferase (NanoLuc) complementation assay. In this assay, the Nbs were fused to LgBiT and their recruitment to μOR measured by protein complementation with plasma membrane-tethered Lyn-SmBiT. Nb63 produced significantly higher recruitment signals than Nb33 across all DAMGO concentrations (Fig. 1H), consistent with stronger μOR binding. TIR-FM assays confirmed enhanced μOR recruitment of Nb63 relative to Nb33 when both probes were co-expressed in the same cell (Fig. 1I).

Finally, we tested Nb63 binding to the other three OR family members, given that the known (Nb33) and predicted (Nb63) Nb binding interfaces are highly conserved (Suppl. Fig. S2A). Nb63 was rapidly recruited to agonist-activated δOR, κOR, and NOP-R (Suppl. Fig. S2B-D), with all showing significantly enhanced recruitment relative to Nb33 (Fig. 1J).

In sum, by changing four CDR2 residues, we obtained Nb63, a variant that binds more strongly to all OR family members compared to the original OR Nbs. Unlike Nb62, which harbors additional framework mutations, Nb63 shows fast OR engagement, making it a promising candidate for effectively blocking OR signaling.

### Intracellular nanobodies block transducer coupling and opioid receptor internalization

We next asked whether the high-affinity variant Nb63 blocks the engagement of intracellular signaling partners. As a proxy for G protein engagement, we performed split-NanoLuc complementation assays in HEK-μOR cells expressing LgBiT-miniGi and Lyn-SmBiT. Nb33 expression had no effect on miniGi recruitment to μOR, while Nb63, expressed at comparable levels (Suppl. Fig. S3A), markedly reduced miniGi association with μOR across all DAMGO concentrations (Fig. 2A).

**Figure 2.**
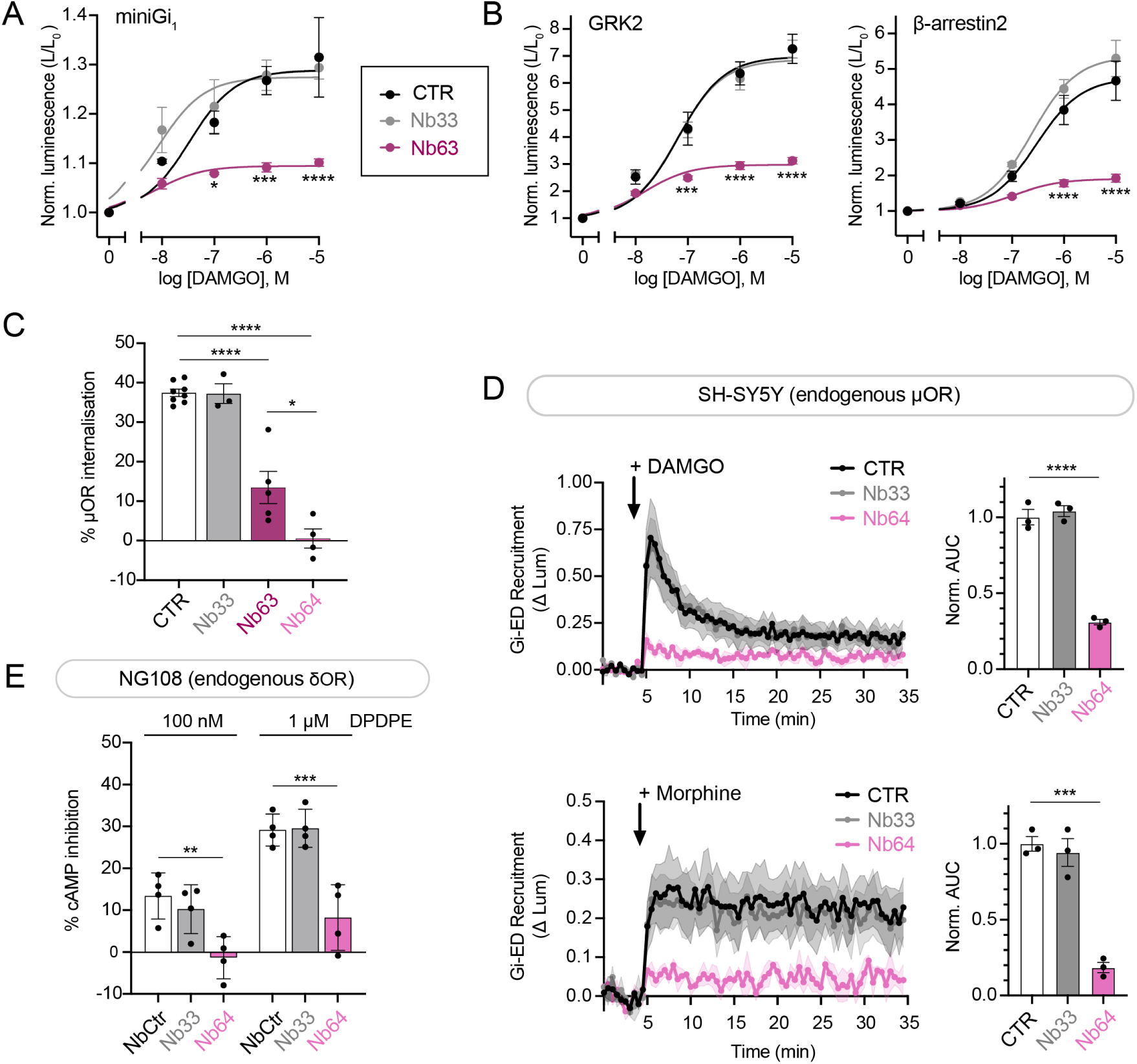
Nanobodies block OR coupling to transducers, receptor internalization, and signal transduction. (A,B) DAMGO concentration-dependent recruitment of transducer proteins to μOR in HEK293 cells expressing EGFP-Nb33, EGFP-Nb63, or EGFP (CTR) as measured by NanoLuc complementation. Data are normalized to mock-treated controls, mean +/- SEM. Regression curves with Hill slope of 1 are shown. Statistical analysis: two-way ANOVA with Dunnett’s multiple comparisons test (*P < 0.05, ***P < 0.001, ****P < 0.0001). (A) Recruitment of miniGi1-LgBiT to plasma membrane-localized Lyn-SmBiT upon μOR activation in stable HEK-μOR cells. N = 3. (B) Recruitment of GRK2-LgBiT (N = 3) or β-arrestin2-LgBiT (N = 5) to μOR-SmBiT in transfected HEK293 cells. (C) DAMGO-induced μOR internalization in HEK-μOR cells expressing EGFP (CTR), EGFP-Nb33, EGFP-Nb63, or EGFP-Nb64 following treatment with 10 µM DAMGO for 25 min. Surface FLAG-μOR was stained with anti-FLAG M1-AF647 and quantified by flow cytometry. Surface μOR levels were normalized to cells not exposed to DAMGO. N = 3 to 8. Statistical analysis: one-way ANOVA with Tukey’s multiple comparisons (*P < 0.05, ****P < 0.0001). (D) Gi/o activity upon μOR activation in SH-SY5Y cells, measured by recruitment of a SmBiT-fused Gi/o effector domain (GI-ED) to the plasma membrane labeled with LgBiT-CAAX following addition of 1 µM DAMGO or 1 µM morphine. Cells were transfected with EGFP (CTR), EGFP-Nb33, or EGFP-Nb64. Left: Quantification of kinetic data via area under the curve (AUC). N = 3, error bars represent +/- SEM. Statistical analysis: one-way ANOVA with Tukey’s multiple comparisons (****P < 0.0001). (E) cAMP response in NG108 cells, measured using the luminescence-based GloSensor assay. Cells were stimulated with forskolin (FSK, 2.5 μM) and inhibitory effect of δOR agonist DPDPE (100 nM or 1 μM) determined. Cells were transfected with non-targeting EGFP-CtrNb (CtrNb is NbALFA ^65^), EGFP-Nb33, or EGFP-Nb64. N = 4, error bars represent +/- SEM. Statistical analysis: one-way ANOVA with Tukey’s multiple comparisons (*P < 0.05, **P < 0.01).

To determine whether Nb63 also inhibits the binding of other transducer proteins, we measured the binding of LgBiT-tagged GRK2 or β-arrestin2 to μOR-SmBiT. In Nb63-expressing cells, the recruitment of GRK2 and β-arrestin2 was strongly blocked, while Nb33 expression again showed no effect (Fig. 2B). The results indicate that Nb63 effectively prevents transducer interactions through high-affinity binding to activated receptors.

Because transducer coupling promotes subsequent receptor endocytosis, we also examined whether Nb63 inhibits agonist-induced μOR internalization. HEK-μOR cells expressing EGFP-tagged Nbs were stimulated with DAMGO and surface receptors were labeled with AF647-conjugated anti-FLAG antibodies prior to quantification by flow cytometry (Suppl. Fig. S3B). Among cells gated for equivalent Nb expression levels (Suppl. Fig. S3B), Nb33 expression did not affect μOR internalization, whereas Nb63 strongly prevented it (Fig. 2C). The findings confirm that intracellular Nb63 uncouples μOR from signaling and downstream regulation.

To maximally suppress residual transducer engagement, we introduced a single framework substitution (A75R) predicted to strengthen μOR contacts without compromising kinetics (Suppl. Fig. S3C). TIR-FM imaging showed that Nb64 retained rapid association and dissociation kinetics (Suppl. Fig. S3D). When intracellularly expressed, Nb64 blocked μOR internalization even more strongly than Nb63, achieving complete inhibition of agonist-driven endocytosis (Fig. 2C), and maintaining pronounced suppression of transducer coupling (Suppl. Fig. S3E). None of the Nbs altered the steady-state μOR surface levels (Suppl. Fig. S3F).

We further compared how intracellular levels of Nb63 and Nb64 influence their ability to inhibit μOR internalization. Nb-expressing cells were grouped into five expression bins based on EGFP fluorescence measured by flow cytometry (Suppl. Fig. S3G). Both Nbs strongly inhibited μOR internalization at intermediate to high expression levels, with Nb64 producing significant inhibition already at lower intracellular abundance (Suppl. Fig. S3H). Thus, Nb64 is better suited than Nb63 to blocking OR signaling over a broad intracellular concentration range and was therefore selected for use in all subsequent functional studies.

### Nb64 inhibits endogenous opioid receptor signaling in neuronal cell lines

Since our goal was to develop an Nb-based tool capable of inhibiting endogenous OR signaling, we next examined Nb64’s function in SH-SY5Y neuroblastoma cells, which endogenously express μORs ^29^. Following μOR stimulation, we monitored Gi/o activity in intact cells by monitoring the recruitment of a Gi/o effector domain (Gi-ED) to the plasma membrane using a previously established NanoLuc assay ^30,31^. The activation of endogenous μORs by either DAMGO or morphine yielded markedly diminished Gi/o activation in Nb64-expressing cells, while Nb33 expression had no effect (Fig. 2D). The findings demonstrate that Nb64 effectively blocks the effects of both peptide and small molecule agonists in a cellular model expressing native μOR.

To assess whether Nb64 also inhibits other members of the OR family, we next examined NG108 cells, which endogenously express functional δORs ^32^. We quantified the Gi/o-mediated inhibition of cyclic AMP (cAMP) accumulation upon δOR activation by DPDPE using the genetically encoded cAMP GloSensor. In Nb64 expressing cells, the δOR-driven inhibition of the cAMP response was markedly reduced relative to control cells expressing a non-targeting control nanobody (NbCtr), while Nb33 expression had no impact on δOR signaling (Fig. 2E).

Overall, these experiments demonstrate that Nb64 is sufficiently potent to override physiological coupling to downstream partners and blunt signaling from endogenous opioid receptor systems.

### Using Nb64 to tune compartment-specific opioid receptor signaling

Having established that Nb64 can robustly control native OR signaling, we next asked whether this inhibition could be spatially confined within the cell. To understand how compartmentalized GPCR signaling determines the physiological responses to endogenous ligands and therapeutic drugs, the development of tools that can modulate GPCR with subcellular precision is a key prerequisite ^3,33,34^. We tested whether Nb64 could be engineered for site-specific OR inhibition.

We focused on the Golgi organelle, since δORs reside in the Golgi at steady state and undergo local activation upon stimulation by membrane-permeant small-molecule agonists ^11,23,25,35^. To target Nb64 to the Golgi, we fused it to the N-terminus of giantin’s Golgi-localization domain and tagged it with HA to monitor subcellular localization (Fig. 3A). In HeLa cells, Nb64-giantin (termed Golgi-Nb64) localized exclusively to the Golgi, as confirmed by co-localization with the Golgi marker ManII (Fig. 3B).

**Figure 3.**
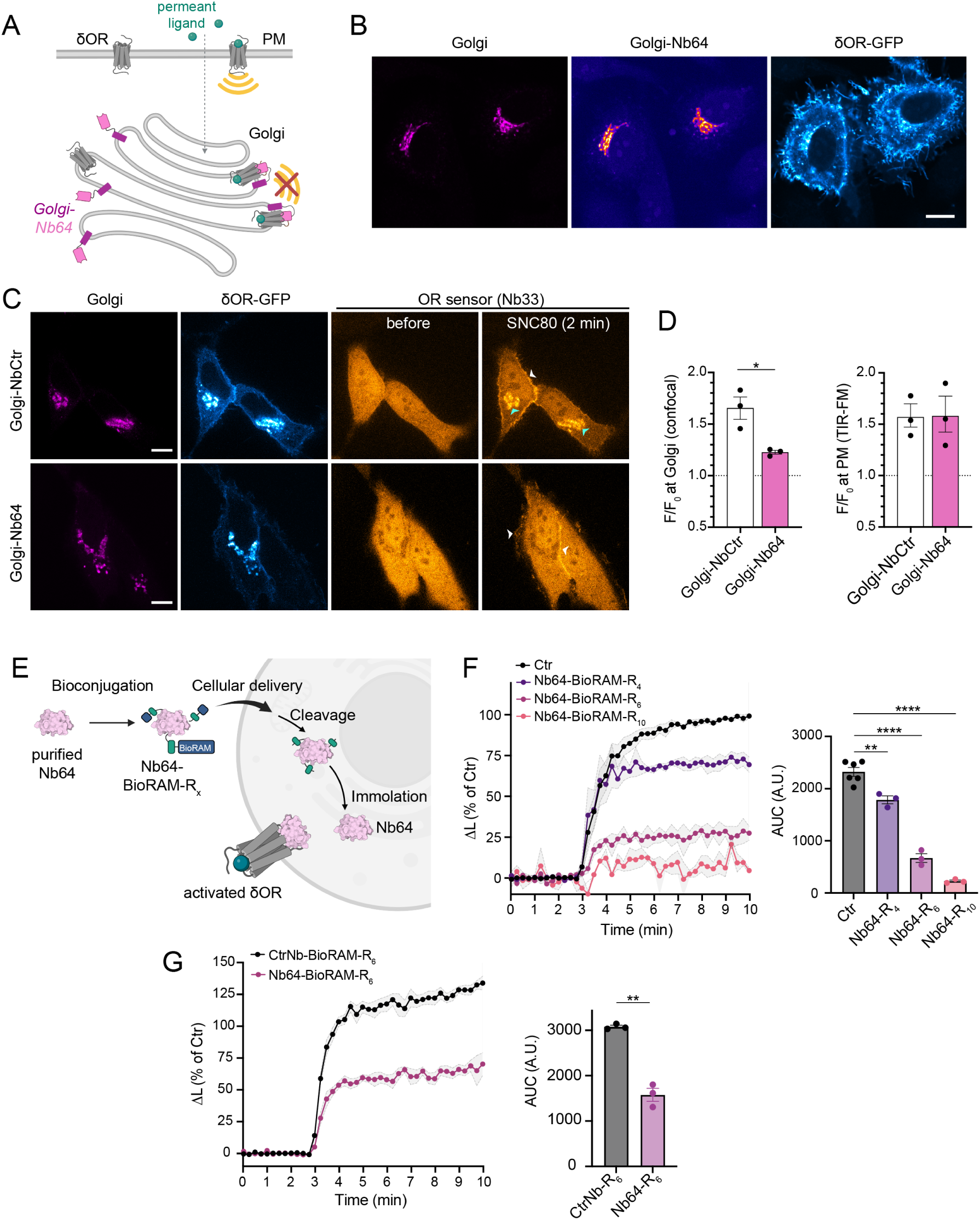
Subcellular targeting and reversible CPP-conjugation expand the functional applications of Nb64. (A) Schematic illustrating Golgi targeting of Nb64 and the resulting inhibition of Golgi-localized δOR in response to membrane-permeant opioid agonists. (B) Confocal images of HeLa cells expressing giantin-HA-Nb64 (Golgi-Nb64, stained with antiHA antibody), GFP-δOR, and the Golgi marker BFP-ManII. Scale bar: 10 μm. (C) Confocal images of HeLa cells expressing GFP-δOR, ManII-BFP, and the OR activation biosensor pmApple-Nb33, together with either Golgi-NbCtr (Golgi-NbALFA, top) or Golgi-Nb64 (bottom), before and 2 min after addition of 1 μM SNC80. Blue arrowheads indicate δOR activation in the Golgi; white arrowheads indicate activation at the plasma membrane. Scale bars: 10 μm. (D) Quantification of OR biosensor recruitment to δOR upon addition of 1 μM SNC80. Left: recruitment to Golgi-localized δOR measured in ManII-positive regions by confocal microscopy. Right: recruitment to plasma membrane δOR measured by TIR-FM. N = 3 (>20 cells), mean +/- SEM. Statistical analysis: two-tailed unpaired t-test (*P < 0.05). (E) Schematic of the BioReversible Arginine Modification (BioRAM) strategy for Nb delivery: (i) selective conjugation of CPPs to aliphatic amines on Nb64, (ii) cellular delivery, (iii) intracellular reduction of the disulfide linkage releasing CPPs, and (iv) traceless self-immolation of the linker to regenerate native Nb64, which then blocks activated ORs. (F) Split NanoLuc miniGi recruitment assay in cells expressing δOR-SmBiT and miniGi-LgBiT. Cells were incubated for 1 h with 5 μM Nb64-BioRAM-Rx conjugates or vehicle in serum-free medium. Luminescence values were normalized to baseline and to vehicle-treated controls (0 = pre-DPDPE; 100 = maximal post-DPDPE). Right: Area under the curve (AUC) analysis. N = 3, mean +/- SEM. Statistical analysis: one-way ANOVA with Tukey’s multiple comparisons test (**P < 0.01; ***P < 0.001). (G) Split NanoLuc miniGi recruitment assay following 1 h incubation with 5 μM Nb64-BioRAM-R6 or CtrNb-BioRAM-R6 in the presence of 10% serum (CtrNb is GBP1 ^66^). Luminescence normalization and AUC analysis as in (F). N = 3, mean +/- SEM. Statistical analysis: two-tailed paired t-test (**P < 0.01).

When we treated cells expressing GFP-tagged δOR and a Golgi-localized NbCtr, the addition of the permeant ligand SNC80 drove strong recruitment of the OR activation biosensor Nb33 to plasma membrane- and Golgi-localized δOR, consistent with previous findings (Fig. 3C,D). In contrast, Golgi-targeted Nb64 strongly inhibited biosensor recruitment to Golgi-δOR, while leaving plasma membrane δOR activation intact (Fig. 3C,D). NanoLuc recruitment assays further showed that miniGi engagement with δOR at the Golgi was significantly reduced in cells expressing Golgi-targeted Nb64 compared to NbCtr (Suppl. Fig. S4A,B).

The data show that Nb64 can be fused to subcellular targeting motifs for site-selective OR inhibition, providing a tool design strategy that can contribute to resolving the physiological role of subcellular OR pools in the future.

### Reversible CPP-conjugation converts Nb64 into an extracellular tool for OR inhibition

We then evaluated whether Nb64 could function without genetic expression, a prerequisite for translational or non-genetic experimental applications. Because Nb64 must access the intracellular face of ORs to block signaling, we rendered purified Nb64 membrane-permeable by conjugating short polyarginine cell-penetrating peptides (CPPs). For this, we employed the recently developed BioRAM (BioReversible Arginine Modification) strategy ^36^, which allowed conjugation of multiple CPPs onto the surface-exposed aliphatic amines (lysine residues and the N-terminus) (Suppl. Fig. S5A). Once internalized, the reductive environment of the cytoplasm reduces the disulfide-linked CPPs, followed by self-immolation of the linker, yielding native Nb64 (Fig. 3E).

To identify optimal Nb64-BioRAM variants, we generated conjugates using CPPs of different polyarginine lengths (BioRAM-R4, -R6, and -R10). Efficient CPP attachment was confirmed by high resolution mass spectrometry and SDS-PAGE, both showing the expected mass shifts toward higher molecular weight (Suppl. Fig. S5B-F). Recombinantly expressed Nb64 contains four potential conjugating sites (Lys43, Lys86, Lys131 as well as the N-terminus). Reducing conditions efficiently removed CPPs, yielding an Nb64 species indistinguishable from the unmodified Nb (Suppl. Fig. S5G), consistent with complete, traceless linker cleavage.

We then assessed whether extracellularly applied Nb64-BioRAMs inhibit miniGi engagement with activated δOR using the NanoLuc complementation assay. All three conjugates significantly reduced miniGi recruitment relative to non-treated controls, with BioRAM-R6 and -R10 showing more pronounced inhibition (Fig. 3F). Previous work established that cell exposure to 5 µM BioRAM-R4 and -R6 conjugates is free of cytotoxicity, whereas BioRAM-R10 conjugates can show cytotoxicity likely due to nonspecific membrane rupture ^36^. Consistent with these prior findings, cells treated with Nb64-BioRAM-R10 displayed a strong reduction in baseline luminescence relative to control, R4, or R6 conditions (Suppl. Fig. S5H), suggesting unwanted cellular effects. Based on this profile, we selected Nb64-BioRAM-R6 for further study.

To test serum compatibility, we applied Nb64-BioRAM-R6 or a non-targeting CtrNb-BioRAM-R6 extracellularly in the presence of 10% serum. Nb64-BioRAM-R6 significantly reduced agonist-driven miniGi recruitment to δOR relative to the control conjugate (Fig. 3G, Suppl. Fig. S5I).

Together, the results show that Nb64 retains its inhibitory function when applied extracellularly as a BioRAM-conjugated, cell-permeant Nb, extending its utility beyond genetic expression and enabling rapid, reversible inhibition of OR signaling in settings where genetic manipulation is impractical or undesirable.

### Nb64 expression in VTA GABAergic neurons in mice inhibits fentanyl-induced dopamine transients and locomotion

We next examined whether Nb64 inhibits μOR signal transduction *in vivo* within the mesolimbic circuit connecting the ventral tegmental area (VTA) to the nucleus accumbens (NAc), a pathway central to opioid reward ^37,38^. Opioid drugs increase dopamine (DA) release in the NAc by disinhibiting VTA DA neurons, which are under tonic suppression by μOR-expressing GABAergic interneurons ^8,39,40^. Activation of μORs on these GABA neurons reduces inhibitory tone onto DA neurons, thereby elevating their activity and increasing DA release in the NAc. We hypothesized that expressing Nb64 selectively in VTA GABAergic interneurons would block this μOR-dependent disinhibition and consequently reduce fentanyl-evoked DA release.

To test this, we expressed Nb64 or the non-targeting NbCtr in VTA GABAergic neurons by bilaterally injecting VGAT::Cre mice ^41^ with recombinant AAVs containing a double-floxed inverted open reading frame (DIO) encoding mRuby2-tagged Nb64 or NbCtr (Fig. 4A,B). To monitor DA dynamics, we delivered the genetically encoded DA indicator dLight1.3b ^42^ via AAV injection into the NAc and implanted an optic fiber above the injection site for photometry recordings (Fig. 4A). Histological analyses confirmed robust Nb64-mRuby2 expression in VTA neurons positioned adjacent to tyrosine hydroxylase (TH)-positive dopamine neurons (Fig. 4B) and revealed isolated Nb64-positive projections in proximity to DA neuron somata, consistent with GABAergic contacts (Fig. 4C). Nb64 expression was well tolerated, with clear cytoplasmic mRuby2 labeling in putative GABAergic neurons (Fig. 4D). Expression of dLight1.3b and fiber placement in the NAc was confirmed for both groups (Fig. 4E, Suppl. Fig. S6).

**Figure 4.**
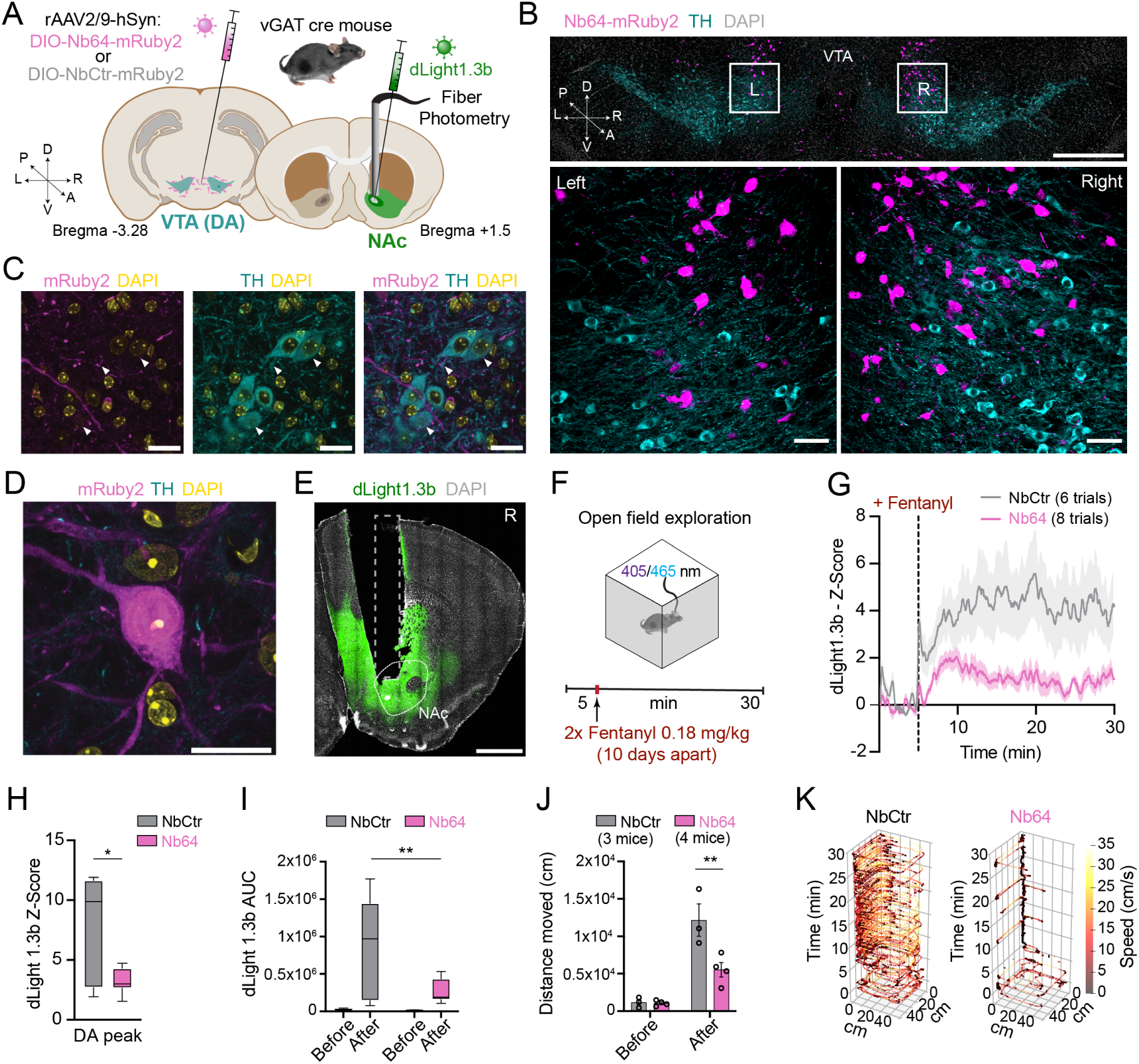
Targeted Nb64 expression in VTA GABAergic neurons blocks fentanyl-induced dopaminergic activation in vivo. (A) Experimental design: expression of Nbs in Ventral Tegmental Area (VTA) GABAergic neurons and dopamine (DA) recordings in the nucleus accumbens (NAc). VGAT::Cre mice were transduced with rAAVs encoding either double-floxed inverted open-reading frame (DIO) hSyn-DIO-Nb64-mRuby2 or hSyn-DIO-NbCtr-mRuby2 (control, NbCtr is NbALFA). The NAc received an rAAV encoding hSyn-dLight1.3b and a fiber optic implant for photometry. (B) Top: Histological confirmation of rAAV-mediated Nb expression (mRuby2, magenta) in VTA GABAergic neurons. A single angled injection achieved bilateral transduction. NomRuby2 expression was detected in dopaminergic neurons (tyrosine hydroxylase (TH), cyan). Scale bar, 500 µm. Bottom: Magnified views of each hemisphere. Scale bar, 50 µm. DAPI (grey) marks nuclei used for tissue registration. (C) Putative GABAergic projections (mRuby2) showing contacts (white arrows) on VTA DA neurons (TH, cyan). DAPI (yellow) marks cell nuclei. (D) Subcellular distribution of Nb64 in VTA neurons. Scale bar, 20 µm. (E) Histological confirmation of dLight1.3b expression in the NAc and fiber optic placement (dotted line square). (F) Experimental setup for fiber photometry recordings in an open field (OF). Fentanyl (0.18 mg/kg) was administered intraperitoneally twice, 10 days apart. (G) 35 min DA fluorescence recording. Fentanyl was injected at minute 5. Data represent mean traces from both sessions (Nb64 group: 8 trials from 4 mice; NbCtr group: 6 trials from 3 mice). (H) Z-score peak after fentanyl injection (*p = 0.0114, unpaired two-tailed t-test). (I) Area under the curve (AUC) of dLight1.3b Z-score comparing before versus after fentanyl injection (**p = 0.0024, two-way ANOVA with Šídák’s multiple comparisons test). (J) Distance traveled (cm) in the OF before versus after fentanyl injection on the first administration day on the first day of fentanyl administration (*p = 0.0031, two-way ANOVA with Šídák’s multiple comparisons test). (K) Tornado plots showing mouse location over time on the first day of fentanyl administration. Color heatmap indicates speed. Left: NbCtr; Right: Nb64.

Because fentanyl increases locomotion in rodents ^8,43^, we performed photometry recordings in an open field to measure DA release and locomotion simultaneously. To ensure that changes in DA levels were driven by fentanyl rather than exploration- or injection-related effects, mice first received saline, which produced neither an increase in NAc DA nor locomotion (Suppl. Fig. S7A). Mice were then administered 0.18 mg/kg fentanyl on two consecutive sessions 10 days apart (Fig. 4F). Fentanyl produced a rapid DA peak (∼30 s after injection) followed by a sustained increase in NAc DA in both groups (Fig. 4G), but the amplitude of the initial peak and the overall DA signal were significantly reduced in Nb64-expressing mice compared to NbCtr controls (Fig. 4H,I). These results demonstrate that Nb64 effectively inhibits fentanyl-driven μOR signaling *in vivo*.

Locomotor behavior, quantified using DeepLabCut ^44^, showed a similar pattern: mice expressing Nb64 moved less than control mice upon first fentanyl injection (Fig. 4J). Furthermore, mice expressing Nb64 moved more slowly (Suppl. Fig. S7B), a difference evident in tornado plots of representative trajectories (Fig. 4K). Together, these findings indicate that Nb64 expression in VTA GABAergic neurons disrupts key neurochemical and behavioral responses to fentanyl, preventing some of the most immediate and physiologically relevant consequences of μOR activation in awake, behaving animals.

## Discussion

Our study establishes intracellular nanobodies as effective tools for controlling OR signaling at molecular, subcellular, cellular, and neural-circuit levels. By combining structure-guided affinity maturation with systematic in-cell functional characterization, we generated Nb64, a high-affinity, active-state-selective inhibitor of ORs that robustly blocks transducer engagement, receptor internalization, and downstream signaling of both endogenous and overexpressed receptors. More broadly, Nb64 defines a mechanistically distinct mode of GPCR inhibition based on intracellular, activation-dependent competitive binding at the transducer interface, rather than ligand engagement at the extracellular receptor surface.

A key strength of Nb64 is the ease of its deployment: once produced in target cells, it resides in the cytosol and can readily access the intracellular face of ORs upon agonist activation, while leaving unoccupied, inactive receptors unchanged. In contrast to orthosteric antagonists or allosteric modulators, which continuously occupy receptors and globally reshape receptor equilibria ^2,45,46^, Nb64 engages ORs selectively and transiently, only after agonist-induced activation has occurred. This activation-dependent mechanism preserves normal receptor organization until signaling is initiated. As a result, Nb64 allows dissection of signaling consequences downstream of receptor activation without altering ligand access, receptor abundance, or basal trafficking dynamics. This mode of action contrasts with biased agonism, which redistributes signaling output by stabilizing specific receptor conformations ^47,48^, and instead enables simultaneous suppression of multiple transducer pathways by steric occlusion at a shared intracellular interface.

The simplicity of genetically encoded nanobodies allows for precise, modular targeting within cells. Being fully protein-based, they can be directed to specific intracellular locations ^3,49^. We demonstrate this by engineering a Golgi-restricted Nb64 that inhibits OR-transducer coupling at this organelle without affecting plasma membrane signaling. Such compartment-specific control is useful for studying spatially segregated GPCR signaling and could be applied to presynaptic terminals, postsynaptic densities, or other specialized neuronal regions ^50,51^. Importantly, this strategy decouples spatial specificity from ligand diffusion, receptor accessibility, and local drug concentration gradients. Instead, inhibition is defined by nanobody localization and stoichiometry, enabling compartment-restricted modulation of GPCR signaling that is difficult to achieve with small-molecule pharmacology.

Photocaged or activatable ligands, while capable of localized action, require complex synthesis, can have limited access to intracellular receptor pools, and involve more elaborate experimental setups ^16,52^. By contrast, intracellular nanobodies can directly interact with native receptors in multiple compartments, allowing studies of localized GPCR signaling that were previously challenging. In addition to spatial specificity, Nb64 can also be adapted for temporal control in the future, e.g. by using inducible expression systems, degradation-based switches, or light-responsive modules, to enable reversible or externally triggered inhibition of OR signaling ^53–56^. Together, these features position intracellular nanobodies as programmable signaling modulators whose activity can be defined by protein engineering rather than ligand chemistry.

Beyond genetic expression, we show that BioRAM-conjugated cell-permeant Nb64 achieves efficient intracellular delivery of functional nanobodies into non-genetically modified cells. This approach can extend nanobody-based modulation to physiologically relevant tissues where transgenic expression is not feasible ^36,57^. Compared with small-molecule antagonists, which often exhibit short half-lives and diffuse broadly, intracellularly delivered Nbs are retained within cells after uptake, providing the potential for sustained inhibition. At the same time, the efficiency, tissue penetration, and temporal resolution of non-genetic delivery of Nb64 will require further optimization for systemic in vivo applications, highlighting complementary roles for genetic and chemical delivery strategies. Nevertheless, the ability to access intracellular receptor interfaces without genetic manipulation establishes a foundation for chemical biology approaches to intracellular GPCR control that extend beyond traditional ligand design.

While nanobodies have been used as extracellular GPCR ligands and hold promise as therapeutic agents for neurological disorders ^58–61^, the potential of intracellular nanobody modulators to study neuromodulation and drug action has remained largely unexplored. Our results demonstrate this potential and suggest that similar strategies could be applied to other intracellular GPCR-targeting nanobodies. More generally, intracellular competitive binders may represent a new class of chemical probes capable of intercepting signaling downstream of receptor activation while preserving endogenous ligand-receptor interactions.

Our in vivo findings demonstrate the functional relevance of Nb64 as a tool for systems neuroscience. Expression of Nb64 in VTA GABAergic neurons significantly attenuated fentanyl-evoked dopamine release in the nucleus accumbens and reduced opioid-driven locomotor behavior, illustrating that intracellular nanobodies can modulate circuit-level outputs by selectively intercepting OR signaling in defined neuronal populations. These experiments establish a proof of principle that cell-type-restricted inhibition of endogenous GPCR signaling is sufficient to reshape neuromodulator dynamics and behavior in vivo.

Looking forward, selective manipulation of other neuronal populations, for example in the pre-Bötzinger complex, could allow Nb64 to dissect the cellular and circuit mechanisms underlying opioid-induced respiratory depression ^62–64^, a major unresolved challenge in opioid pharmacology. More generally, the ability to intercept GPCR signaling downstream of receptor activation provides a route to dissect context-dependent signaling across diverse neuromodulatory, inflammatory, and metabolic systems.

In conclusion, our work establishes Nb64 as a versatile molecular tool that directly connects receptor-level mechanisms with neuronal circuit function. It demonstrates that intracellular nanobodies can be engineered for high-affinity, rapid, activation-dependent, and compartment-specific inhibition of native GPCRs, enabling precise interrogation of neuromodulatory signaling. By implementing intracellular, activation-dependent competitive inhibition of native GPCRs, this study defines a chemical biology strategy for spatially and cell-specifically controlling GPCR signaling within intact biological systems.

## Acknowledgements

We thank all members of the Stoeber lab for constructive feedback and ideas. We thank Jonathan Franke for synthesizing and characterizing the BioRAM reagents used in this study and Raquel Mendes for sharing secondary antibodies. This work was supported by the Swiss National Science Foundation (PCEFP3_181282 to MS, 320030E_224301 and 320030-236030 to TP) and the U.S. National Institutes of Health (R61DA051531 to AM and MvZ). We also acknowledge funding from the European Research Council (ERC) under the European Union’s Horizon 2020 research and innovation program (grant 891959 to TP), the Israel Science Foundation (ISF grants 197/25 and 3960/25 to MK), and support from the EMBO Young Investigator Program (MS and TP).

## Author contributions

ZV designed and functionally characterized intracellular Nbs. PLM co-designed and performed in vivo experiments, with histology analyses by AC, and locomotion analysis by GH. AK conducted the Nb affinity maturation campaign. SP, ARM, and NMF designed Nb variants and performed functional studies. JVA and CPRH designed and conducted the intracellular Nb delivery experiments. ZST performed interaction and structure analyses. MvZ and MK designed and supervised experiments. MS, AM, and TP conceived and directed the study. All authors contributed to data analysis and interpretation. MS wrote the manuscript with input from all authors.

## Materials and methods

### Nanobody affinity maturation

The overall strategy to improve Nb39 affinity was guided by a prior affinity maturation campaign for Nb6B9, which binds to the β2-adrenergic receptor and was matured from Nb80 ^67^. Guided by the structure of mouse μOR bound to Nb39 (PDB: 5C1M) ^24^, 15 sites on Nb39 that formed significant interactions with the intracellular face of the μOR were selected for mutagenesis with designed amino acid substitutions as outlined in Suppl. Fig. S1. The resulting sequences were designed via degenerate primer library design following the method outlined in ^67^. In total, this yielded a nanobody library with theoretical diversity of ∼1.5 x10^9, which was assembled by PCR and introduced into S. cerevisiae strain EBY100 using a standard lithium chloride-based electroporation protocol with the yeast display vector pYDS1.0. The transfected library had an estimated ∼1.0 x 10^9 transformants based on dilution plating onto SDCAA medium. To enrich for high affinity variants, we carried out rounds of positive selection with decreasing amounts of BU72-bound μOR, which was purified as described previously ^24^. We found that attempts to negatively select against μOR bound to the inverse agonist BNTX or neutral antagonist naloxone led to depletion of active-state binding clones, as these ligands do not fully inhibit the receptor’s ability to sample active conformations. Instead, we performed three rounds of positive selection for BU72-bound μOR using magnetic separation columns (MACS, Miltenyi) using Anti-Alexa Fluor 647 magnetic beads and anti-M1 FLAG-Alexa Fluor 647 antibody. To deplete Nbs that might bind the antibody reagent, each round of selection was first cleared against M1 FLAG-Alexa Fluor 647 antibody and Anti-Alexa Fluor 647 magnetic beads. To identify slowly dissociating clones, we performed a kinetic selection procedure, accomplished by first incubating the library of remaining clones after positive selection with BU72-bound μOR, followed by extensive washing of unbound receptor, and finally, incubation with a 100x molar excess of soluble Nb39. In this setup, the soluble Nb39 competitively prevents receptors that have unbound from library clones from rebinding. After examining the kinetics of μOR unbinding from the bulk library, we positively selected clones still bound to receptor after 3 hours of incubation at room temperature with saturating excess Nb39 using MACS after an initial clear against anti-Flag M1-AF647 antibody. Finally, to enrich the highest affinity clones, we performed one round of FACS-based selection against receptors bound to BU72. At this stage, a random sequencing of 20 clones yielded 7 unique sequences (each with multiple repetitions). Yeast expressing each of these seven clones were then assayed individually in dose response with increasing concentration of BU72-bound μOR to determine a pseudo-affinity; this analysis revealed Nb62 as a particularly high affinity clone.

### Protein structure analyses

The following structures of complexes of active opioid receptor with nanobody 39 (Nb39) were used in our analyses (with PDB IDs): μOR with Nb39 (5C1M) ^24^ and two structures of κOR with Nb39 (6B73, 7YIT) ^68,69^. Three missing residues (259-261) in KOR (6B73) were modelled with the homology modeling program Nest (Xiang et al., 2002) using the structure of the MOR in PDB ID 5C1M as a template. The following missing residues were modelled *ab-initio* with Loopy ^70^: residues 103-109 in Nb39 (5C1M), Nb39 residues 103-106 in chain B and residues 102-106 in chain D (6B73), and Nb39 residues 103-109 and KOR residues 259-261 (7YIT). Missing atoms and side chains were predicted using Scap ^70^. Hydrogen atoms were added using CHARMM, and the structures were subjected to conjugate gradient minimization with a harmonic restraint force of 50 kcal/mol/Å^2^ applied to the heavy atoms. 3D structural visualisations were carried out with the PyMol molecular graphics program (https://www.pymol.org/).

### Mammalian cell culture conditions

HEK293 cells (CRL-1573, ATCC, female) or HeLa cells (CRM-CCL-2, ATCC, female) were cultured in Dulbecco’s modified Eagle’s medium (DMEM, Gibco), supplemented with 10% fetal bovine serum (FBS, GIBCO). HEK293 cells stably expressing N-terminally FLAG-tagged μOR (HEK-μOR) were cultured in the presence of 250 μg/ml Geneticin (Gibco). NG108-15 cells were cultured in DMEM supplemented with 10% FBS, 0.1 mM hypoxanthine, 400 nM aminopterin, 0.016 mM thymidine and 1.5 g/L sodium bicarbonate. For transient DNA expression, Lipofectamine 2000 (Invitrogen) was used according to the manufacturer’s instructions. The full sequence of all plasmids used in this study will be made accessible via the Yareta data repository.

### Ligands

DAMGO [D-Ala2, N-Me-Phe4, Gly5-ol]-enkephalin acetate salt (catalog no. E7384), DPDPE [D-Pen2,5]-enkephalin hydrate (catalog no. E3888), and morphine sulfate (catalog no. 1448005) were purchased from Sigma-Aldrich. SNC80 (catalog no. 0764), Nociceptin (NOP, catalog no. 0910/1), Dynorphin A (DynA, catalog no. 3195), ICI 174,864 (catalog no. 0820) were purchased from Tocris. Fentanyl (CAS no.1443-54-5) was obtained from the University Hospital Zurich with authorization from the Canton of Zurich.

### TIR-FM protein recruitment assay

HEK293 cells and HEK-μOR cells were seeded on poly-L-lysine-coated 35 mm glass-bottomed dishes and transfected with EGFP-Nb33, EGFP-Nb62 (0.3 μg), EGFP-Nb63 or EGFP-64 (0.25 μg) using 3 µL Lipofectamine 2000. 0.8 μg mRuby-miniGi1 (mGi1) was transfected for the mGi1 recruitment assays. HEK293 cells without stable receptor expression were additionally transfected with FLAG-δOR, FLAG-κOR or FLAG-NOP-R (1 μg DNA). 16-24 h after transfection, surface ORs were labeled for 10 min with anti-FLAG M1-AF647 and media changed to HBS imaging solution (Hepes buffered saline with 135 mM NaCl, 5 mM KCl, 0.4 mM MgCl2,1.8 mM CaCl2, 20 mM Hepes, 1 mM d-glucose, 1% FBS, adjusted to pH 7.4). Cells were imaged at 37°C using a Nikon Eclipse Ti microscope using a 100x 1.49 Oil CFI Apochromat TIR-FM objective, temperature chamber, objective heater, perfect focus system and an ORCA-Fusion BT Digital CMOS camera. Live cell image series were acquired every 5 s before and 5 min after agonist addition. Protein relocalization (dF) was calculated as F(t)/F0 with F(t) = Nb or mGi1 signal after ligand addition and F0 = Nb or mGi1 signal before ligand addition.

### Split NanoLuc-based Nb and transducer recruitment assay

HEK293 cells were seeded into 6-well plates and individual wells transfected with 0.25 μg of i) Lyn-SmBiT and LgBiT-Nb, ii) Lyn-SmBiT and LgBiT-miniGi1 ^11^, iii) μOR-SmBiT and GRK2-LgBiT or iv) μOR-SmBiT and LgBiT-β-arrestin2, in combination with 0.3 μg EGFP or EGFP-Nb33 or 0.25 μg EGFP-Nb63 or EGFP-Nb64, using 3 µL Lipofectamine 2000. Binding of Nb, miniGi1, GRK2 or β-arrestin2 to ORs triggers NanoLuc (LgBiT and SmBiT) complementation. 16-24 h after transfection, cells were seeded into black, clear-bottom 384-well plates (20,000 cells/well) in FluoroBrite DMEM (Gibco) and the NanoLuc substrate Nano-Glo (Promega, N2012), and incubated for 45 min at 37°C. The luminescence signal was recorded using the FDSS/μCELL kinetic plate imager (Hamamatsu) with an integrated simultaneous dispensing head and simultaneous detection across the plate. After acquiring baseline luminescence for 3 min, agonists (concentration specified in figure legends) were added to the cells. Luminescence was recorded every 2 s for 7 min post agonist addition. Luminescence values were normalized to vehicle treated control cells and baseline signal (before agonist).

### Western Blot to probe EGFP levels

To compare EGFP levels, 24 h after transfection, cells were lysed in RIPA buffer (10 mM Tris pH 7.4, 1X Triton, 150 mM NaCl, 0.1% SDS, 2 mM EDTA) supplemented with protease and phosphatase inhibitors (Roche) and lysates were sonicated followed by protein quantification using the Pierce BCA protein assay (Thermo Fisher). Protein samples were mixed with NuPAGE LDS sample buffer and 100 mM DTT, and heated for 10 min at 70°C. Equal quantities of lysates were separated by SDS-PAGE using 4-12% BisTris Plus gels (Thermo Fisher) and transferred to an isopropanol-activated PVDF membrane. After blocking in TBS with 5% BSA, membranes were incubated overnight at 4°C with anti-EGFP and anti-α-tubulin primary antibodies (1:2000) in TBS-Tween (0.1% v/v) with 5% BSA. After washing, HRP-coupled anti-mouse (1:10000) and anti-rabbit (1:4000) secondary antibodies were added for 1 h at RT. Membranes were imaged in the presence of Pierce ECL SuperSignal West Pico plus (Thermo Fisher) using the iBright 1500 device (Invitrogen).

### Flow cytometry-based μOR internalization assay

To quantify μOR internalization, 16-24 h after transfection, HEK-μOR cells were treated with DAMGO (10 μM) or vehicle control for 25 mins. Cells were immediately washed in ice-cold PBS and surface μOR were labelled for 20 min with anti-FLAG M1-AF647 in ice-cold PBS while undergoing shaking to detach cells. Cells were resuspended and analyzed by flow cytometry using the Beckman Coulter Cytoflex.

### Split NanoLuc-based Gi activation assay in SHSY5Y cells

SH-SY5Y cells were plated in 6-well dishes and transfected with Lipofectamine 2000 24 hours before the assay (1 µg Gi-ED-SmBiT, 200 ng LgBiT-CAAX, and 200 ng EGFP or EGFP-Nb64). On the day of the assay, cells were lifted using TrypLE (Gibco 12604021) and centrifuged at 500xg for 3 minutes. Cells were resuspended in assay buffer (in mM: 135 NaCl, 5 KCl, 20 HEPES, 5 D-glucose, 1.8 CaCl2, and 0.4 MgCl2, pH 7.4) supplemented with 5 µM coelenterazine-H (Research Products International). Cells were plated into white 96-well plates (Corning 3912) in a volume of 100µL (100,000 cells/well). DAMGO and morphine were diluted in assay buffer with 5 µM coelenterazine-H. Prior to the start of the assay, the plate was equilibrated for 10 min in a Tecan Spark platereader warmed to 37 °C. Luminescence values were acquired at 30 sec intervals with an integration time of 500 ms. After a 5 min baseline, vehicle or drug was added and luminescence was recorded for an additional 30 minutes. For each well, luminescence was first normalized to the average luminescence during the baseline period. The average normalized luminescence for vehicle-treated wells was then subtracted from the drug-treated wells at each time point. Each condition was performed as 3 technical replicates, which were subsequently averaged.

### Luminescence-based live cell cAMP accumulation assay in NG108 cells

A 10 cm dish of NG108 cells was transiently transfected with GloSensor-20F cAMP reporter (2.5 μg, Promega), and EGFP, EGFP-Nb33, (2.1 μg) or EGFP-Nb64 (1.6 μg). 24 h post transfection, cells were plated in 96-well dishes and incubated in FluoroBrite DMEM (Gibco) containing 250 μg/ml luciferin. After acquiring baseline luminescence for 3 min, Forskolin (2.5 μM) and varying concentrations of DPDPE were added to the cells. Luminescence was recorded every 2 s for 7 min post Forskolin addition. Measurements were collected every 5 s for 30 min using a HAMAMATSU FDSS microCELL platereader and luminescence values were normalized to vehicle-treated control cells and baseline signal (before Forskolin).

### Confocal microscopy-based Nb33 recruitment assay

HeLa cells were seeded on poly-L-lysine-coated 35 mm glass-bottomed dishes. After 24 h, cells were transfected with combinations of SEP-δOR (0.8 μg DNA), ManII-BFP (0.2 μg), pmApple-Nb33 (0.3 μg), Nb64-giantin, CtrNb-giantin (ALFANb) using 3 μL Lipofectamine 2000. Cells were live-imaged 16-24 h post transfection in HBS imaging solution. Recruitment of pmApple-Nb33 was imaged with a Spinning disk confocal microscope (Nipkow, Zeiss) using an EC Plan Neofluar 100x/1.3 Oil Ph3 objective in a temperature and CO2-controlled environment (37°C, 5% CO2). Images of the same cells were acquired before and 3 min after 100 nM SNC80 addition. Nb recruitment to Golgi-localised ORs was quantified as the agonist-induced fold-change of signal in the Golgi area (ManII-defined mask) normalised to cytosol area relative to before agonist addition.

### Split NanoLuc-based miniGi1 recruitment assay

Hela cells were seeded into 6-well plates and transfected with 0.4μg of δOR-SmBiT and 0.25μg LgBiT-miniGi1, in combination with i) 0.25μg EGFP, ii) 0.25μg EGFP-Nb64, iii) 0.6 μg Golgi-Nb64 (GFP-TGN-Nb64), iv) 0.6μg Golgi-CtrNb (GFP-TGN-NbALFA) using 3μL Lipofectamine 2000) per well. 16-24 h after transfection, cells were seeded into black, clear bottom 384-well plates (20,000 cells/well) in FluoroBrite DMEM (Gibco) and the NanoLuc substrate Nano-Glo (Promega, N2012), and incubated for 45 min at 37°C. The luminescence signal was recorded using the FDSS/μCELL kinetic plate imager (Hamamatsu) with an integrated dispensing head and simultaneous detection across the plate. After acquiring baseline luminescence for 3 min, agonists were added to the cells. Luminescence was recorded every 1.5 s for 7 min post agonist addition. A combination of the peptide antagonist ICI 174,864 (100 μM, added 5 min before baseline acquisition) and 100 nM SNC80 was used to block δOR at the PM and activate only Golgi-localized receptors. Luminescence values were normalized to vehicle treated control cells and baseline signal (before agonist addition). Recruitment kinetics are represented as the percentage of the maximal luminescence signal (10 max values) detected for each agonist. Area under the curve (AUC) analysis was performed using the data normalized to maximum (% of max).

### Split NanoLuc-based miniGi1 recruitment assay to evaluate Nb64-BioRAM conjugates

The Split Nanoluc complementation assay protocol was adapted to measure the miniGi1 recruitment to δOR in the presence of BioRAM-conjugated Nbs. HeLa cells were transfected with 0.4μg of δOR-SmBiT/0.25μg LgBiT-miniGi1, as described above and seeded in 384-well plates (20,000 cells/well) in FluoroBrite DMEM and Nano-Glo substrate, in the presence or absence of 10% FBS. The cells were incubated for 1h at 37°C with the Nb64-BioRAM conjugates (BioRAM4, BioRAM6, BioRAM10) or NbCtr-BioRAM6 at 5 μM concentration. Luminescence was recorded before and after agonist addition every 1.5 sec, as described above. Luminescence values were normalized to vehicle-treated control cells and baseline signal for each BioRAM conjugate, and the average of 10 measurements was used as a single time point, to smoothen the curves. Recruitment kinetics are shown as the percentage of the maximal luminescence signal (5max values) detected in the absence of BioRAM conjugates. AUC was calculated from the data normalized to maximum (% of max).

### Nb64 expression, purification, and characterization

Nb64 was expressed in WK6 E. coli and purified upon periplasmic extractions similar to NbE described in ^59^. In brief, Nb64 was isolated by His-tag affinity purification. Ni-NTA beads (Thermo) were equilibrated with wash buffer (50 mM Tris pH 7.5, 150 mM NaCl, 10 mM imidazole). Periplasmic extracts were added into a column of pre-equilibriated Ni-NTA beads and allowed to elute by gravity. The eluate was added for the second time to beads to ensure complete retrieval of His-Nb64. The beads were washed with 10 column volumes (CV) of wash buffer, followed by the addition of 5 CV of elution buffer (50 mM Tris pH 7.5, 150 mM NaCl, 500 mM imidazole). The eluted fractions with the highest concentration of protein were collected, followed by one-pot TEV-cleavage and dialysis against the storage buffer (20 mM Hepes pH 7.4, 150 mM NaCl, 10% glycerol). Dialyzed protein fractions were concentrated using Amicon spin filtration (Amicon® Ultra-0.5, 3-10k MWCO) to a final concentration of 100 µM. Protein concentration was determined using a Nanodrop (Nanodrop One/OneC, Thermo Scientific, USA). Samples were flash-frozen in liquid nitrogen and stored at -80°C. Protein conjugates were characterized by SDS-PAGE and high-resolution mass spectroscopy (HR-MS). Briefly, HR-MS spectra were recorded on a Waters H-class instrument equipped with a quaternary solvent manager, a Waters sample manager-FTN, a Waters PDA detector and a Waters column manager with an Acquity UPLC protein BEH C18 column (1.7 μm, 2.1 mm x 50 mm). Samples were eluted with a flow rate of 0.3 mL/min. The following gradient was used: “QTof”: 0.01% FA in H2O; B: 0.01% FA in MeCN. 5 % B: 0-1 min; 5 to 95 % B: 1-7min; 95 % B: 7 to 8.5 min. Mass analysis was conducted with a Waters XEVO G2-XS QTof analyzer.

### BioRAM conjugation and characterization

BioRAM reagents were chemically synthesized according to ^36^. Recombinantly expressed Nb64 (50 µM) was alkylated by adding TCEP (2.5 eq) and iodoacetamide (10 eq) for 1 h at 25°C. Excess small molecules were removed and the protein was buffer exchanged HEPES buffer (200 mM HEPES at pH = 8.25, 140 mM NaCl) using Zeba Spin purification columns. BioRAM reagents (20 eq) were added to the buffer exchanged Nb64. The reaction mixture was incubated at 25 °C for 3 hours at 500 rpm shaking. The reaction mixture developed a yellow color due to the nitrophenol byproduct generated during successful protein conjugation as well as from gradual hydrolysis over time. The resulting modified protein was separated from excess peptide by using an Amicon spin filtration (Amicon Ultra-0.5, 3-10k MWCO) until a 1:100 dilution, followed by desalted and rebuffered by Zeba Spin purification in storage buffer (20 mM HEPES at pH = 7.5, 150 mM NaCl, 10% glycerol). Protein concentration was determined using a Nanodrop (Nanodrop One/OneC, Thermo Scientific, USA). The samples were flash-frozen in liquid nitrogen and stored at -80°C. Protein conjugates were characterized by SDS-PAGE and HR-MS as mentioned above. As CtrNb, the Green fluorescent protein binding protein 1 (GBP1) was used ^66^. Expression, purification, and BioRAM conjugation of GBP1 was similar to Nb64.

### Animals

Animal procedures were performed in accordance with the guidelines of the Animal Welfare Ordinance (TSchV 455.1) of the Swiss Federal Food Safety and Veterinary Office and were approved by the Zürich Cantonal Veterinary Office. VGAT:mice (B6J.129S6(FVB)-Slc32a1tm2(cre)Lowl/MwarJ) of both sexes and 3-4 months of age were used. Mice were housed in a temperature- and humidity-controlled environment on a normal 12-hour light-dark cycle, with food and water available ad libitum.

### Animal surgeries and viral injections

For in vivo fiber photometry experiments during open field exploration, stereotactic surgeries were performed on isoflurane-anesthetized VGAT:cre mice. After removing the scalp, two craniotomies were performed to deliver the recombinant adeno-associated viruses (rAAVs) through a glass micro-pipette controlled by a micro manipulator (NanoJect III). AAV9 encoding either Nb64 (∼1.0 × 10^13^ GC/mL) or NbCtr (NbALFA) (∼1.1 × 10^13^ GC/mL) were injected in the VTA. For specific targeting GABAergic neurons, these constructs included a double inverted open frame (DIO) and both were under the hSyn promoter. A single 10° angle injection (400 nL) was performed to target VTA on both hemispheres: (-3.28 mm AP, +0.9 mm ML, 4.3 mm DV). A separate rAAV encoding dLight1.3b (∼7.9 × 10^12^ GC/mL) was injected (400 nL) into the right nucleus accumbens (NAc): (+1.5 mm AP, +0.7 mm ML, - 4.5 mm DV). An optic fiber was implanted 0.2 mm above the NAc injection site (400 μm core diameter, NA = 0.57, Neurophotometrics) on the same surgery day. Mice recovered for at least 3 weeks before behavioral experiments.

### Photometry recording and analysis

Fiber photometry recordings in the NAc were performed using an iFMC6 photometry system equipped with excitation filters E1 (460–490 nm) and E2 (555–570 nm), emission filters F1 (500–540 nm) and F2 (580–680 nm), an isosbestic channel IE (400–410 nm), and a short-pass filter S was used, controlled by the Doric Neuroscience Studio v6.1.2.0 software. A low-autofluorescence patch cord (400 μm, 0.57 N.A., Doric Lenses) was attached to the metallic ferrule on the mouse head for photometry recordings. A 465 nm LED was used to excite dLight1.3b (∼30 µW at the patchcord tip) while emitted light was measured by a photodiode detector (Newport). A 405 nm LED was used to excite the indicator at the isoemissive wavelength and the resulting fluorescence was used as the control fluorescence signal. Signals were sinusoidally modulated at 208 Hz and 572 Hz (405 nm and 465 nm, respectively) via lock-in amplification, then demodulated on-line and low-passed filtered at 10 Hz. Mice were connected to the patchcord 5 min before open field (OF) exploration in a new cage. To do this, mice were placed in the center of the OF arena (50 x 50 cm) for 35 minutes while dLight1.3b transients were being recorded. Mice were picked up from the OF at min 4:45 and injected at ∼5:00 intraperitoneally. On day 1, mice were injected with saline, day 3 with fentanyl (0.18 mg/kg) and on day 13 again with fentanyl (0.18 mg/kg). Analysis of the raw photometry data was performed using a custom-written MATLAB script ^44^. Firstly, all raw signals were denoised with a low-pass Butterworth filter (of 10 Hz cutoff frequency) to remove high-frequency noise components. Next, the denoised fluorescent signals were corrected for photobleaching and motion artifacts. For bleaching correction, a single polynomial fit was used (only to the first 5 min of recording) for each signal independently which was then detrended from the filtered traces. Motion correction was performed by finding the best linear fit of the isoemissive (405 nm) control signal to the 465 nm functional signal. Given that mice were freely moving, the estimated animal motion-dependent component was subtracted from the photobleaching-corrected functional signals ^71^. Z-Score normalization was performed by first calculating the mean of the signal during a 4 min period prior to the injection and subtracting this from the functional signals across the whole recording length. Afterwards, the signals were divided by the standard deviation of the same pre-injection period. Peak z-score values were estimated as the maximum z-score value of the mean signals across the whole recording duration. To assess the cumulative effect of the signal triggered by fentanyl administration, the area under the curve (AUC) was computed using the trapezoidal method (trapz MATLAB function) on the mean trial z-scored values.

### Locomotion behavior analysis during open field exploration

Mice were placed in a 50 x 50 cm OF arena, and their behavior was recorded using a Basler camera (acA640-750um) at 25 frames per second. Raw videos were fed into DeepLabCut (DLC) ^44^ where anchor points on several mouse body parts were manually added. The arena limits were defined by adding anchor points to each corner. The mouse position was determined as the center-body location. Each mouse anchor point coordinate was transformed using the arena corners’ coordinates in each video. DLC processed all videos in an unsupervised manner. Low confidence tracking points (likelihood < 0.6) and outliers were replaced with the previous valid coordinate (pseudo-interpolation). All coordinates’ matrices were truncated to exactly 30 minutes by removing the final ∼5 min to standardize across animals/sessions and divided into pre-injection (5 min) and post-injection (25 min) epochs. Data were temporally downsampled by selecting every 3rd frame (bin size = 3) to reduce computational load while preserving movement patterns. Instantaneous speed was calculated as Euclidean distance between consecutive sampled frames divided by time interval (0.12 sec). Velocity values were smoothed using a 3-frame rolling average. Total distance travelled was computed as the sum of frame-to-frame displacements. Tornado plots were generated to show the mouse location (x/y), and speed (color map) over the course of the experiments (z). A custom code was written on Python (version 3.11.5) using the following libraries: OpenCV (homography transformation), NumPy (numerical computations), pandas (data manipulation), matplotlib (visualization).

### Histology and confocal Imaging

Brains were sectioned in 50 µm thick coronal slices using a vibratome (VT1200S, Leica Biosystems) and collected in 1x PBS. To confirm the fiber location and expression of dLight1.3, two sections were selected within a 150 µm range around the fiber in the NAc. In addition, three sections in the range of 250 µm around the VTA were chosen to assess overall transduction of midbrain GABAergic neurons with the Nb64 or NbCtr expression (VTA overview). In a second experiment, other two sections were chosen from the VTA for higher magnification images to ascertain that Nb64 and NbCtr were well expressed in cellular compartments (e.g., in somata). First, sections were blocked for 2 h at room temperature in a solution containing 5% BSA and 0.3% Triton in PBS. The dLight1.3 expression was visualized by immunolabeling the GFP on the indicator (chicken anti-GFP, 1:1000; catalog no. GFP-1010, Aves Labs). The VTA sections for the overview were incubated with a primary antibody against tyrosine hydroxylase (TH, rabbit anti-TH, 1:3000; catalog nr. AB152, Merck Millipore) and tdTomato (goat anti-tdTomato, 1:1000, catalog nr. TA150129, OriGene Technologies), which also recognizes mRuby2 fused to the Nbs. For Nb images, the sections were incubated with the same primary antibodies as listed above. However, a higher concentration of the TH AB was used (1:2000). All primary antibodies were incubated at 4°C overnight. The sections were washed (3x 10 min, 1x PBS), followed by secondary antibody incubation at room temperature for 2 h. The NAc sections were incubated with Alexa Fluor 488 donkey anti-chicken (1:500; catalog no. 703-545-155, Jackson laboratories), while the VTA overview sections were incubated with Alexa Fluor 488 goat anti-rabbit (1:500; catalog nr. 111-545-003, Jackson laboratories) and Alexa Fluor 568 donkey anti-goat (1:1000; catalog nr. A-11057, Invitrogen). The Nb image sections were incubated with Alexa Fluor 647 donkey anti-rabbit (1:500; catalog nr. 711-607-003, Jackson ImmunoResearch) and the same secondary antibody to stain against mRuby2 as for the VTA overview images. Then, the sections were washed again (2x 10min, 1x PBS) and incubated with 4′,6-diamidino-2-phenylindole (DAPI, diluted in blocking solution with factor 1:10’000, 1x 10 min) before being mounted on microscope slides using Hydromount (catalog no. HS-106, National Diagnostics). All images were taken at a confocal laser scanning microscope (Axio Imager LSM 800, Zeiss). The images of the NAc and the VTA overview images were acquired with a 25x oil-immersion objective (i LCI Plan-Neofluar 25x/0.8 Imm Korr DIC M27, Zeiss) with 0.5x digital zoom. The tile format was set to 512x512 pixels per tile before stitching and four z-stacks were imaged. A bit depth of 8 was selected for these images. The Nb images were acquired using a 63x oil-immersion objective (Plan-Apochromat 63x/1.40 Oil DIC M27, Zeiss) with 1x or 1.6x digital zoom. They are a single tile of 1024x1024 pixels and a z-step size of 0.71 µm. A bit depth of 16 was selected. The Smart Setup function of the ZEN microscopy software (version 2.6, blue edition) was used to select the optimal acquisition settings for the fluorophores to be imaged.

