## Supplementary Figures for "An engineered nanobody inhibitor for molecular-to-circuit control of opioid receptor function"

#### Supplementary Figure S1

A

```
                 FR1                CDR1               FR2       CDR2
      1  .........10........20........30........40........50........60
Nb33  1  MAQVQLVESGGGLVRPGGSRRLSCVDSERTSYPMGWFRRAPGKEREFVASITWSGIDPTY
Nb39  1  MAQVQLVESGGGLVRPGGSLRLSCVDSERTSYPMGWFRRAPGKEREFVASITWSGIDPTY
Nb62  1  MAQVQLVESGGGLVRPGGSLRLSCVDSERTSYPMGWFRRAPGKEREFVASISWDGIFPEY
Nb63  1  MAQVQLVESGGGLVRPGGSRRLSCVDSERTSYPMGWFRRAPGKEREFVASISWDGIFPEY

                 FR3                         CDR3
     61  .........70........80........90........100.......110.......120.....
Nb33 61  ADSVADRFTISRDVANNTLYLQMNSLKHEDTAVYYCAARAPVGQSSSPYDYDYWGQGTQVTVSSA
Nb39 61  ADSVADRFTTSRDVANNTLYLQMNSLKHEDTAVYYCAARAPVGQSSSPYDYDYWGQGTQVTVSSA
Nb62 61  ADSVADRFTTSRDAYNNTLYLQMNSLKHEDTAVYYCAARAPVGQSSSPYDYDYWGQGTQVTVSSA
Nb63 61  ADSVADRFTISRDVANNTLYLQMNSLKHEDTAVYYCAARAPVGQSSSPYDYDYWGQGTQVTVSSA
```

B

| Position | 51 | 52 | 53 | 54 | 55 | 56 | 57 | 58 | 59 | 65 | 69 | 71 | 73 | 74 | 75 | On-yeast Affinity | |
|---|---|---|---|---|---|---|---|---|---|---|---|---|---|---|---|---|---|
| Nb39 | I | T | W | S | G | I | D | P | T | A | T | S | D | V | A | 1.5 uM | |
| Library | FILV | STAG | WLMR | SGN DTA | GS | FILV +W | DNHY FLIV | PYSH | TNK ADE | ADNT | TNSY | TNSY | DG | VTAI | ADYS | | |
| Clone 1 | I | S | W | D | G | I | F | P | T | A | T | S | D | A | Y | 10 nM | |
| Clone 2 | I | S | W | D | G | I | F | P | E | A | T | S | D | A | Y | 10 nM | = Nb62 |
| Clone 3 | I | T | W | D | G | I | F | P | T | A | T | S | D | A | Y | 15 nM | |
| Clone 4 | I | T | W | D | G | L | F | P | D | A | T | S | D | A | A | 20 nM | |
| Clone 5 | I | T | W | D | G | I | F | P | D | A | T | S | D | V | A | 20 nM | |
| Clone 6 | I | T | W | D | G | L | F | P | D | A | T | S | D | A | Y | 20 nM | |
| Clone 7 | I | T | W | D | G | L | Y | P | D | A | T | S | D | A | Y | 20 nM | |

C

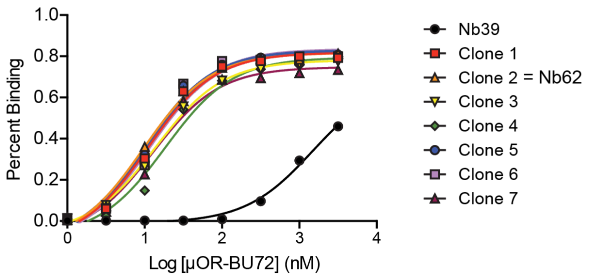


D

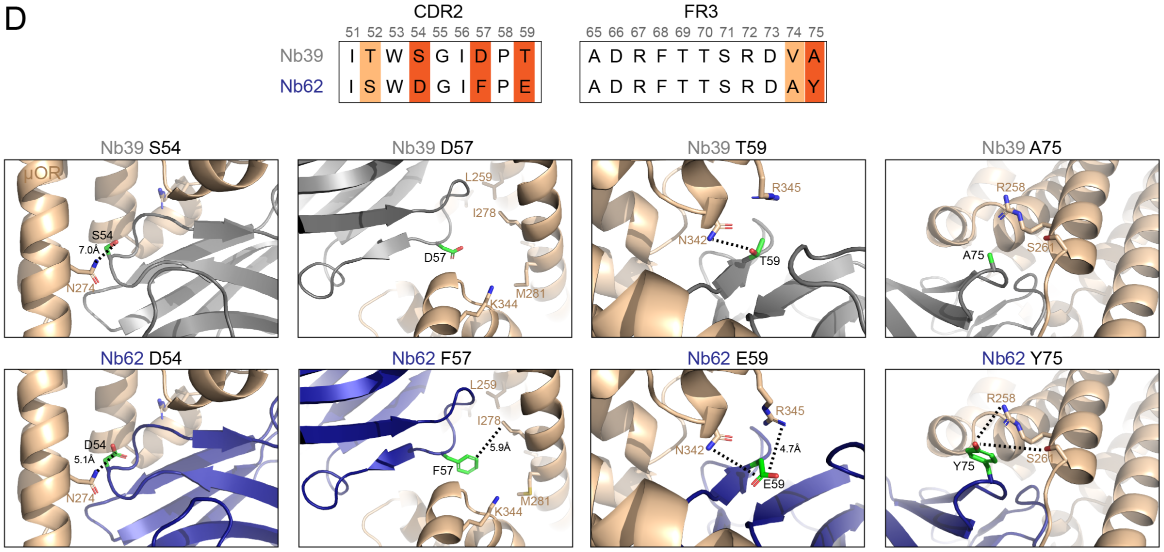


**Supplementary Figure S1. High-affinity nanobodies for the active-state µ-opioid receptor.**

(A) Sequence alignment of the original nanobodies Nb39 and Nb33, and the novel high-affinity variants Nb62 and Nb63.

(B) Residues in the CDR2 and framework region of Nb39 located at the μOR interface were substituted. The amino acid sequence of parental Nb39 is shown in the top row, and the Nb library of possible residue substitutions is shown below. The residues of the seven Nb clones obtained after selection round 5 are shown along with their on-yeast titration EC50 values. Clone 2 corresponds to Nb62.

(C) On-yeast titration of Nb-expressing yeast with BU72-bound μOR-647. All seven high-affinity variants exhibit markedly improved affinity relative to parental Nb39.

(D) Residues in the CDR2 and third framework region of Nb62 that may increase affinity for μOR relative to Nb39: S54D in Nb62 enables a hydrogen bond with N274 of μOR; D57F in Nb62 introduces non-polar interactions with I278 of μOR; T59E in Nb62 enables a salt bridge with R345 of μOR; and A75Y in the third framework region of Nb62 enables two potential hydrogen bonds with S261 or R258 of μOR.

#### Supplementary Figure S2

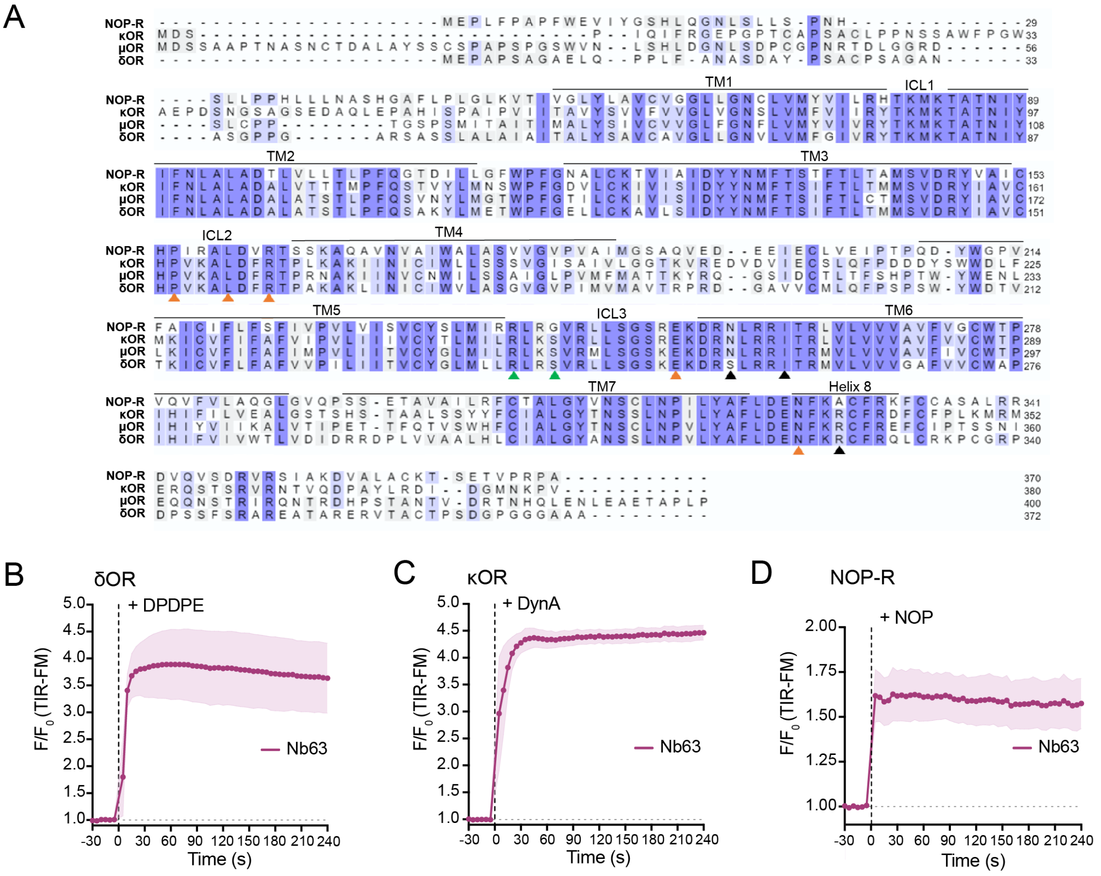


**Supplementary Figure S2. High-affinity nanobodies bind all active-state opioid receptors.**

(A) Multiple sequence alignment of OR family members (mouse) generated with UniProt. Orange arrowheads indicate the Nb39-interacting residues based on the Nb39-μOR crystal structure (PDB: 5C1M). Black arrowheads indicate predicted interactions enabled by mutations in the CDR2 region of Nb62 and Nb63. Green arrowheads indicate predicted interactions enabled by the framework mutation in Nb64 (see Fig. 3). Residues are highlighted according to similarity.

(B–D) Quantification of TIR-FM time-lapse imaging of HEK293 cells expressing EGFP-Nb63. Frames were acquired every 5 s. F0 represents the average fluorescence intensity before agonist addition. Cells co-expressed the indicated opioid receptor and were treated with 10 μM of the corresponding agonist: (B) δOR expression with DPDPE treatment; (C) κOR expression with dynorphin A (1-17, DynA) treatment; (D) NOP-R expression with nociceptin/orphanin FQ (NOP) treatment.

#### Supplementary Figure S3

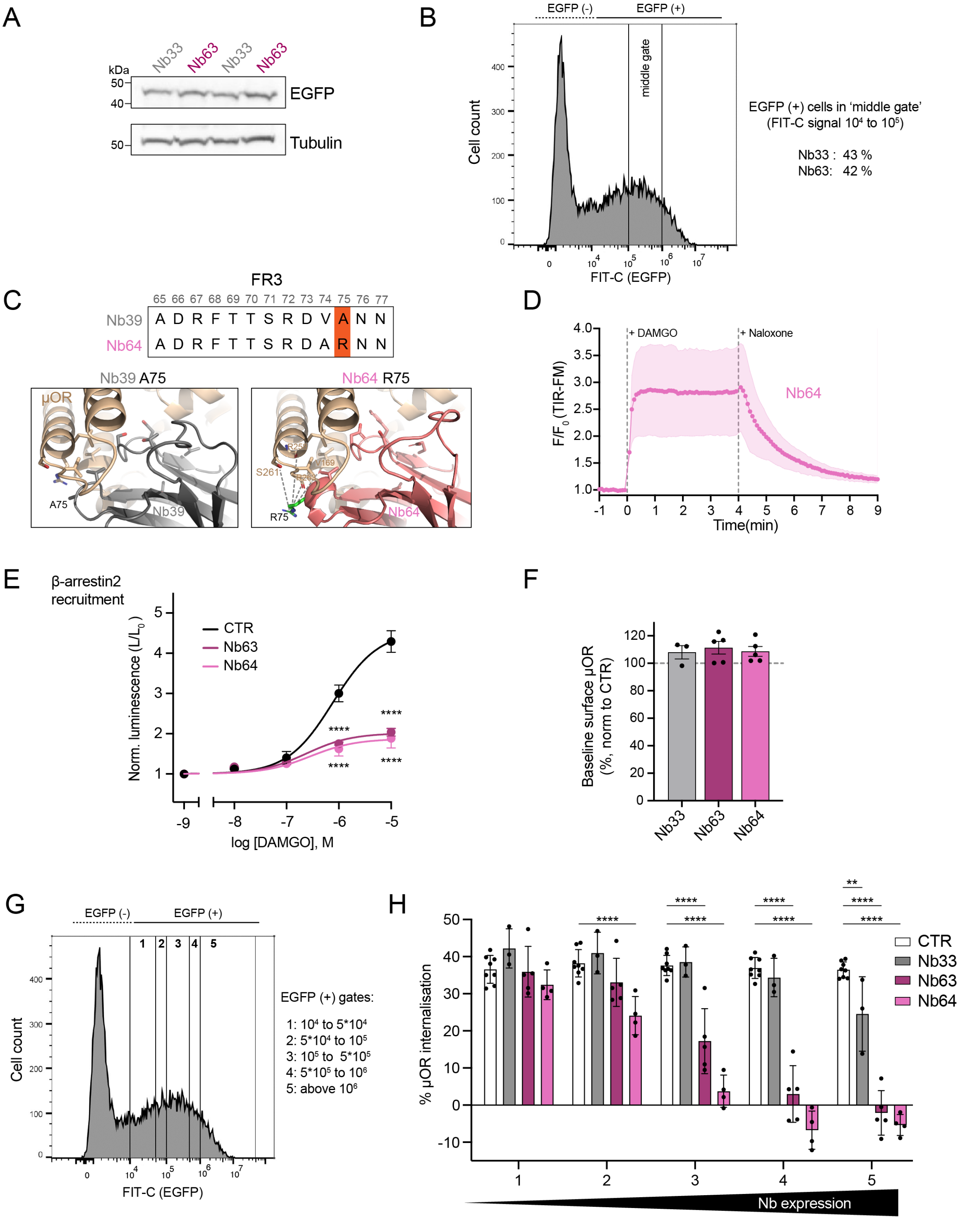


**Supplementary Figure S3. Characterization of Nb63 and Nb64 expression, and functional effects.**

(A) Nb expression levels assessed by Western blot in HEK293 cells transfected with EGFP-Nb33 or EGFP-Nb63. Tubulin served as a loading control. Two independent lysates were loaded per Nb condition.

(B) Flow cytometry histogram showing EGFP-Nb63 expression after gating for live singlet cells. The 'middle gate' indicated here was applied to all conditions of the internalization experiments shown in Fig. 2C.

(C) Predicted interactions of residue R75, introduced in the Nb64 framework, with μOR. R75 may form a hydrogen bond with S261 and additional hydrogen bonds with backbone atoms of R258, R263, or V169, potentially contributing to increased receptor affinity.

(D) Recruitment kinetics of EGFP-Nb64 to μOR in HEK293 cells monitored by TIR-FM time-series (5 s intervals). DAMGO (1 μM) and naloxone (10 μM) were added sequentially. F0 represents the average fluorescence intensity before agonist addition. N = 3, mean +/- SEM.

(E) DAMGO concentration-dependent recruitment of β-arrestin2-LgBiT to μOR-SmBiT in HEK293 cells expressing EGFP (control), EGFP-Nb63, or EGFP-Nb64. Signals were normalized to mock-treated controls. N = 3, mean +/- SEM. Regression curves assume a Hill slope of 1. ****P < 0.0001 by two-way ANOVA with Dunnett's multiple comparisons test.

(F) Surface μOR levels at baseline (no agonist treatment) in HEK-μOR cells expressing EGFP-Nb33, EGFP-Nb63, or EGFP-Nb64. FLAG-tagged μOR was stained with anti-FLAG M1-AF647 and quantified by flow cytometry (gating as in B). Surface μOR levels were normalized to untransfected cells (set to 100%). N = 3 to 5, error bars represent +/- SEM. ns: not significant by one-way ANOVA.

(G) Gates 1 to 5 used to analyze cells with different EGFP-Nb expression levels. Identical gates (shown here based on a flow cytometry histogram of EGFP-Nb63 transfected cells) were applied to all conditions in the internalization data shown in Suppl. Fig. S6H.

(H) DAMGO-induced μOR internalization in cells expressing different Nbs, analyzed across the five EGFP expression bins shown in Fig. S3G. N = 3 to 5, error bars represent +/- SD. *P < 0.05, **P < 0.01, ***P < 0.001, ****P < 0.0001, by two-way ANOVA with Dunnett's multiple comparisons test.

#### Supplementary Figure S4

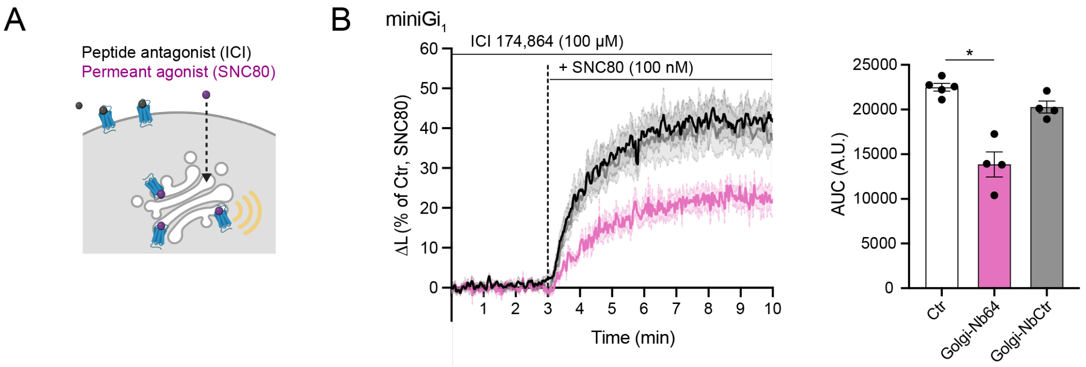


**Supplementary Figure S4. Golgi-Nb64 blocks δOR-mGi coupling at the Golgi apparatus upon permeant opioid drug addition.**

(A) Ligand strategy for selective activation of Golgi-resident δOR (blue), achieved by combining excess membrane-impermeant peptide antagonist (ICI174,864) with the permeant agonist SNC80.

(B) Split NanoLuc miniGi recruitment assay in cells expressing δOR-SmBiT and miniGi-LgBiT together with Golgi-Nb64 (GFP-TGN-Nb64), Golgi-NbCtr (GFP-TGN-NbALFA), or EGFP (Ctr). Cells were pre-incubated for 5 min with ICI174,864 (100 µM) followed by addition of 100 nM SNC80. *Left:* Luminescence traces, normalized to baseline and vehicle-treated controls, then scaled to the SNC80 response (0 = pre-SNC80; 100 = maximal post-SNC80). *Right:* Area under the curve (AUC) quantification. N = 3, mean ± SEM. Statistical significance was assessed using ordinary one-way ANOVA with Tukey's multiple comparisons test (*P < 0.05).

#### Supplementary Figure S5

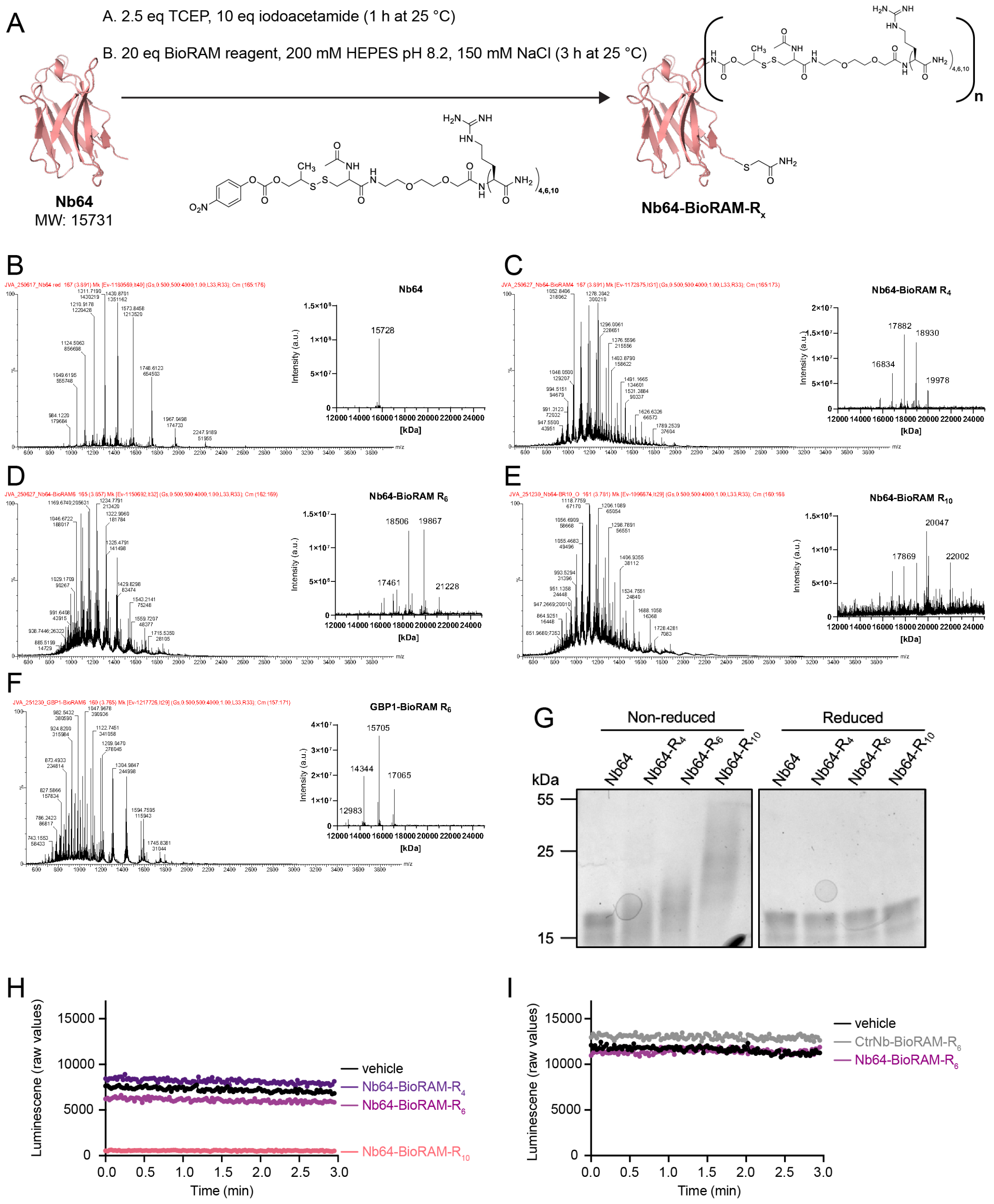


**Supplementary Figure S5. Reversible CPP-conjugation of Nb64 and CtrNb**

(A) Schematic representation of BioRAM conjugation applied to recombinantly expressed Nb64 or non-targeting NbCtr.

(B-F) Characterization of Nb64 and NbCtr-BioRAM bioconjugates using Quadrupole Time-of-Flight high-resolution mass spectrometry (QToF HR-MS). Deconvoluted HR-MS spectra (inset) of (B) Nb64 $[M + H]^+$ calc.: 15731, exp.: 15728, (C) Nb64-BioRAM-$R_4$ $[M + H]^+$ calculated (calc).: 16853, 17900, 18947, 19994; observed (obs.):15788, 16834, 17882, 18930, 19978, (D) Nb64-BioRAM-$R_6$ $[M + H]^+$ calc.: 17162, 18520, 19878, 21236; obs.: 17461, 18506, 19867, 21228, and (E) Nb64-BioRAM-$R_{10}$ [M

+ H]$^+$ calc.: 17921, 20052, 22183; obs.: 17869, 20047, 22002. (F) GBP1 (CtrNb)-BioRAM-$R_6$ [M + H]$^+$ calc.: 12984, 14341, 15699, 17057 ; obs.: 12983, 14344, 15705, 17065.

(G) SDS–PAGE analysis of Nb64 and Nb64-BioRAM conjugates under non-reducing and reducing conditions. Reduction of Nb64-BioRAM conjugates resulted in bands corresponding to the unmodified Nb64, suggesting the recovery of the wild-type Nb64 in reductive conditions.

(H) Baseline luminescence recordings (3 min, raw values) from cells expressing δOR-SmBiT and miniGi-LgBiT after 1 h incubation with 5 μM Nb64-BioRAM-R4, -R6, or -R10.

(I) Baseline luminescence recordings (3 min, raw values) from cells expressing δOR-SmBiT and miniGi-LgBiT after 1 h incubation with 5 μM Nb64-BioRAM-R6 or 5 μM NbCtr-BioRAM-R6.

### Supplementary Figure S6

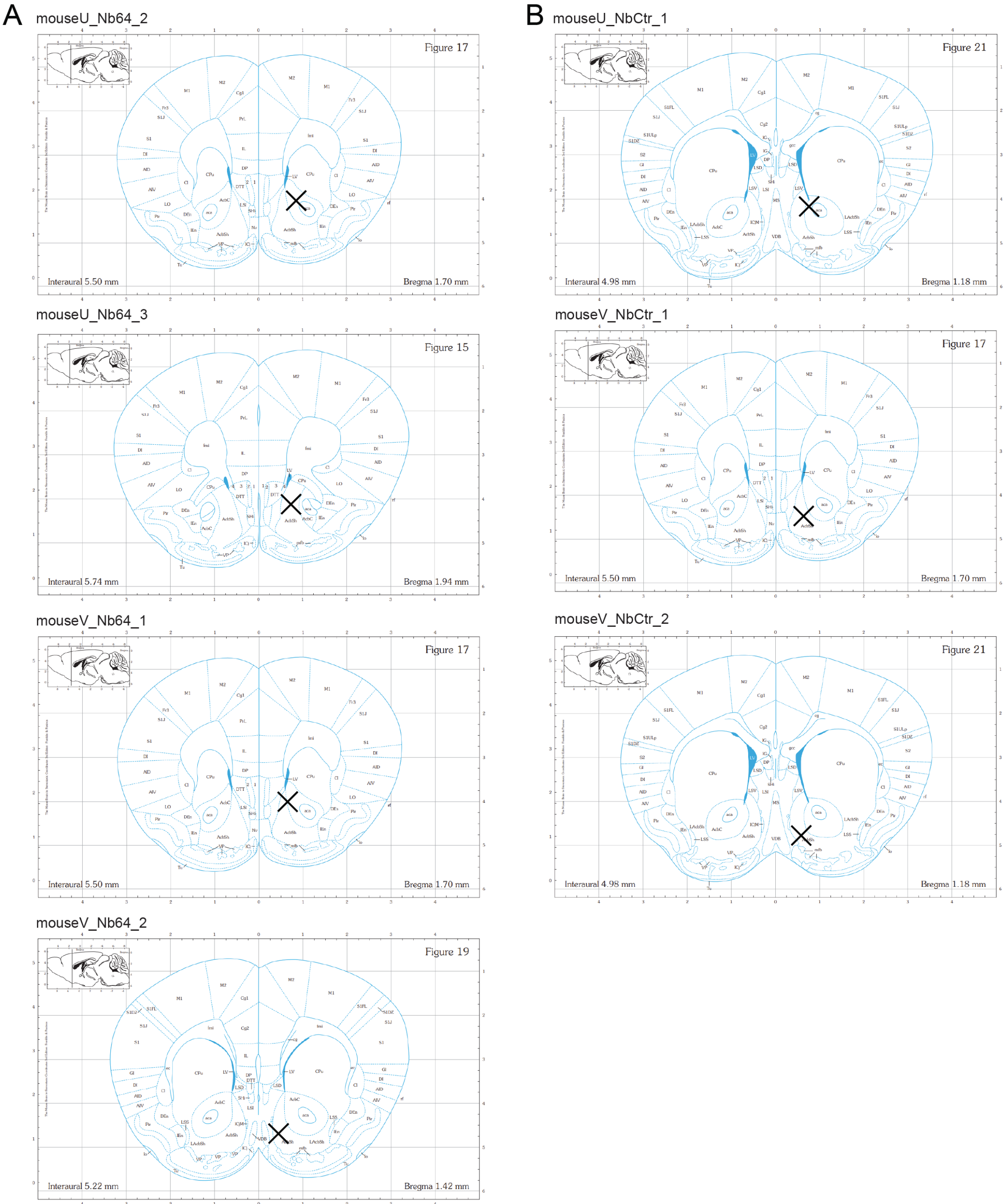


**Supplementary Figure S6. Anatomical location of fiber placements for in vivo photometry recordings.**

(A) Mice from the Nb64 group.
(B) Mice from the NbCtr group.

#### Supplementary Figure S7

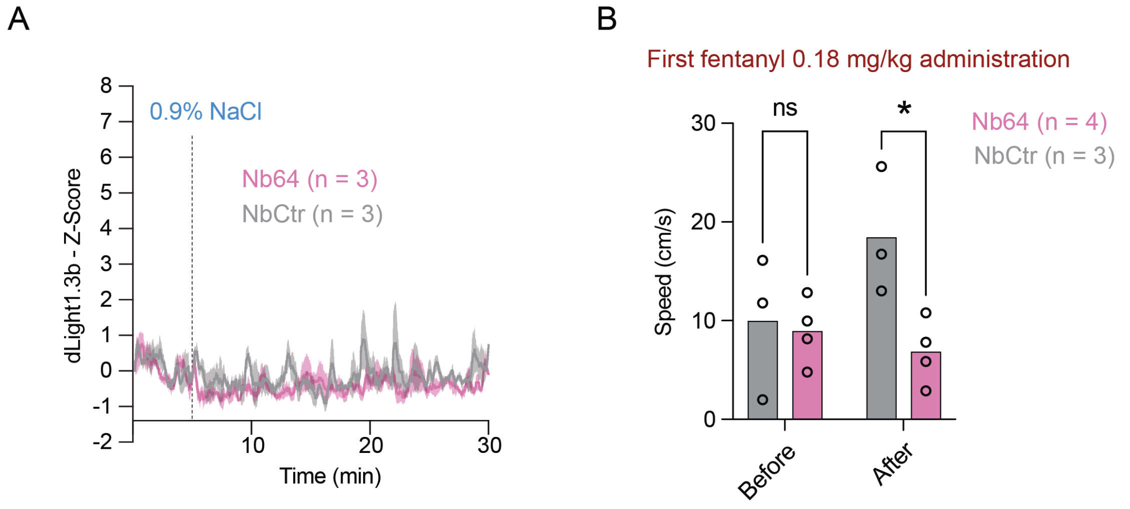


**Supplementary Figure S7. Additional photometry and behavioral analyses of nanobody-expressing mice.**

(A) Intraperitoneal saline injection effect on dopamine (DA) release in nucleus accumbens. Both Nb64 and NbCtr expressions show similar DA transients.
(B) Comparison of speed inside the open field (OF) before vs. after fentanyl injection during the first session. (*p = 0.0263, two-way-ANOVA, Šídák's multiple comparisons test).
